# Basement membrane mechanics drives patterned response to developmental signalling

**DOI:** 10.64898/2026.02.13.705301

**Authors:** Ana Raffaelli, Tom P.J. Wyatt, Claire S. Simon, Léa M.D. Wenger, Kathy K. Niakan, Ewa K. Paluch, Kevin J. Chalut

## Abstract

Precise spatiotemporal patterning of cell fate decisions during development, such as those accompanying gastrulation, has traditionally been attributed to morphogen gradients. Emerging evidence also points to a crucial role for mechanics in regulating cell fate decisions. Here, we uncover that basement membrane mechanics in human pluripotent stem cells (hPSCs) regulates the spatiotemporal response to gastrulation-inducing BMP4. Reducing mechanosensation via soft substrates or chemical inhibitors abolishes stereotypical spatial patterning of the BMP4 response in hPSC colonies. This loss arises from disrupted epithelial polarity and increased permeability, which enhance BMP receptor accessibility. Strikingly, apical exposure to soluble laminin similarly disrupts polarity and suppresses hPSC mechanosensing, phenocopying the soft substrates effect and suggesting feedback between polarity signaling and mechanosensing. Finally, softening the basement membrane in mouse embryos triggers ectopic and premature mesoderm differentiation, disrupting gastrulation patterning. Together, this study describes a mechanochemical feedback mechanism that establishes basement membrane mechanics as a key regulator of developmental patterning.

## INTRODUCTION

Embryonic development relies on successive, precisely patterned fate acquisition events governed by strictly regulated spatiotemporal response to developmental biochemical signals. As such, mechanisms that control the spatial patterning of developmental fate decisions have been extensively studied. Early studies, notably by Alan Turing^1^ and Lewis Wolpert^2^, suggested that both reaction-diffusion processes and morphogen gradients can generate patterns of the developmental signals driving fate acquisition. Since then, developmental patterning processes have been mostly studied through the lens of morphogen gradients and kinetics^3–5^ .

Yet, more recent results suggest that morphogen gradients and kinetics do not fully explain developmental patterning of cell fate. Indeed, studies in several model organisms demonstrate that a uniformly applied biochemical signal can also elicit a spatiotemporally patterned response through patterning of cellular competence, i.e. ability to respond to the signal^6–10^. An example where pattern formation arises from spatial differences in cellular competence is the initiation of gastrulation in the mouse embryo. Gastrulation is triggered by BMP4 signalling, through a feedback loop with Wnt and Nodal, which together drive pluripotent epiblast cells toward mesodermal and endodermal fates ^11–13^. The BMP4 signal is secreted by the extraembryonic ectoderm into the pro-amniotic cavity and is thus apically accessible to most of the epiblast. Yet, gastrulation starts in a spatially restricted manner at the posterior midline of the embryo. Restriction of the BMP response to the posterior epiblast is thought to result from local secretion of BMP inhibitors by the anterior visceral endoderm (AVE)^14,15^. Further, epithelial polarity plays a role by geometrically limiting BMP signalling through BMP receptor sequestration to the basolateral membrane, which renders most epiblast cells incompetent to respond^16^. However, why the posterior midline epiblast cells become competent to respond to BMP4 signal only about a day after BMP4 secretion, and how this response starts in a restricted region at the posterior midline and only then further amplifies during gastrulation, has not been explained.

Modelling of early patterning during human gastrulation has been effectively done using human pluripotent stem cells (hPSCs), which are developmentally similar to human post-implantation epiblast cells^17,18^. In culture, hPSCs form monolayered, epithelialized colonies, and are used to model early fate instruction in human development. For example, it has been shown that upon uniform BMP4 exposure, hPSC colonies undergo radial patterning of fates. In these patterns, ectoderm is in the centre surrounded by, in increasing concentric circles, cells displaying markers of mesoderm, endoderm and extraembryonic-like fates^19,20^ . This fate patterning results from spatial restriction of the response to BMP4 to the colony margin in hPSCs on glass^6,19^. Similar to BMP4 response in mouse gastrulation^16^, the patterned response to BMP4 in hPSCs has been attributed to polarised, basolateral localisation of BMPRs, which would make only the receptors at the colony margin accessible to the BMP4 signal (Extended Data Fig. 1a)^6^. Taken together, previous results suggest that fate patterning in response to BMP4 in pluripotent stem cells, both *in vivo* and *in vitro,* is regulated by patterning of cell competence.

On the other hand, stem cell fate acquisition is known to be influenced by, alongside biochemical signalling, extracellular matrix (ECM) properties such as stiffness. ECM stiffness has been shown to strongly influence cell fate choice in a number of systems^21–29^. In particular, substrate mechanics can modulate mesodermal differentiation of hPSCs in response to mesoderm-inducing signals *in vitro*^30^. In parallel, recent *in vivo* studies have suggested that basement membrane remodelling influences gastrulation, affecting the proportion of differentiating mouse epiblast cells^31,32^. Given these previous observations, we set out to test the hypothesis that mechanical cues from the basement membrane, by regulating cellular competence, drive response to developmental signals to orchestrate cell fate patterning.

To test our hypothesis, we investigate how basement membrane mechanics affects spatiotemporal response to BMP4 in hPSCs. We find that when cultured on soft substrates, hPSCs lose the stereotypical spatially restricted response to BMP4 observed on stiff substrates and instead display response to BMP4 throughout the hPSC colony. Mechanistically, we find that this loss of patterned response stems from a disruption of epithelial polarity due to reduced mechanosensing. Intriguingly, apical laminin delivery to hPSCs on stiff substrates similarly abrogates the spatial restriction of the BMP4 response by disrupting epithelial polarity and mechanosensing. Finally, we demonstrate that softening the basement membrane in the mouse embryo *in vivo* is sufficient to trigger premature and ectopic differentiation towards mesoderm, indicating disruption of spatiotemporal response to developmental signals in the epiblast. Altogether this study provides mechanistic insight into the role of basement membrane mechanics in regulating spatiotemporal response to biochemical signals and thus in patterning fate acquisitions.

## RESULTS

### Soft substrates abrogate the spatial patterning of hPSC response to BMP signalling

To investigate the influence of substrate stiffness on spatiotemporal response of hPSC to mesoderm-inducing BMP4 and subsequent mesodermal differentiation, we cultured hPSCs on polyacrylamide hydrogels^25^ of different stiffnesses (Fig. 1a). We first sought to establish a stiff and a soft substrate condition, on which the cells exhibit the reported overall difference in mesodermal differentiation^30^ but maintain the monolayered epithelial architecture seen in the epiblast. We found that when cultured on hydrogels of stiffnesses between 160 kPa and 3.5 kPa, hPSCs consistently formed monolayers, albeit with different epithelial morphologies. On 3.5 kPa hydrogels, hPSC monolayers exhibited increased height compared to the stiffest hydrogels of 160 kPa (Extended Data Fig. 1b). We further observed that culturing hPSCs on hydrogels of a stiffness lower than 1 kPa induced multilayering and colonies formed domes (Extended Data Fig. 1b), reminiscent of dome-like colonies of naïve pluripotent stem cells. Hence, we selected the 3.5 kPa hydrogels for the soft condition, to ensure that cells remained in an epithelial monolayer as seen at that stage *in vivo*, while the stiffest hydrogels of 160 kPa were used for the stiff condition. Next, we investigated the effect of BMP4 treatment on hPSC differentiation on soft and stiff hydrogels. One of the earliest mesodermal markers upregulated in response to BMP4 is Brachyury, which is encoded by the T-box transcription factor T (*TBXT*)^33^. Therefore, to probe whether substrate stiffness affects BMP4-induced mesoderm differentiation in our experimental system, we assessed expression of Brachyury protein after 24 hours of BMP4 treatment. We found that BMP4 treatment on soft substrates caused a significantly higher level of Brachyury expression in comparison to stiff substrates, both at the protein (Fig. 1b,c) and gene transcript level (Fig. 1d). Soft substrates alone, in the absence of BMP4, were insufficient to cause an increase in expression of *Brachyury* (*TBXT*) (Fig. 1d) or enhance differentiation of hPSCs. Indeed, in the absence of BMP4, hPSCs maintained indistinguishable expression levels of the pluripotency markers *POU5F1, SOX2* and *NANOG* between soft and stiff substrates (Extended Data Fig. 1c). Therefore, we hypothesised that the enhanced mesodermal differentiation observed on soft substrates was caused by a coupling between mechanical and biochemical signalling.

**Figure 1.**
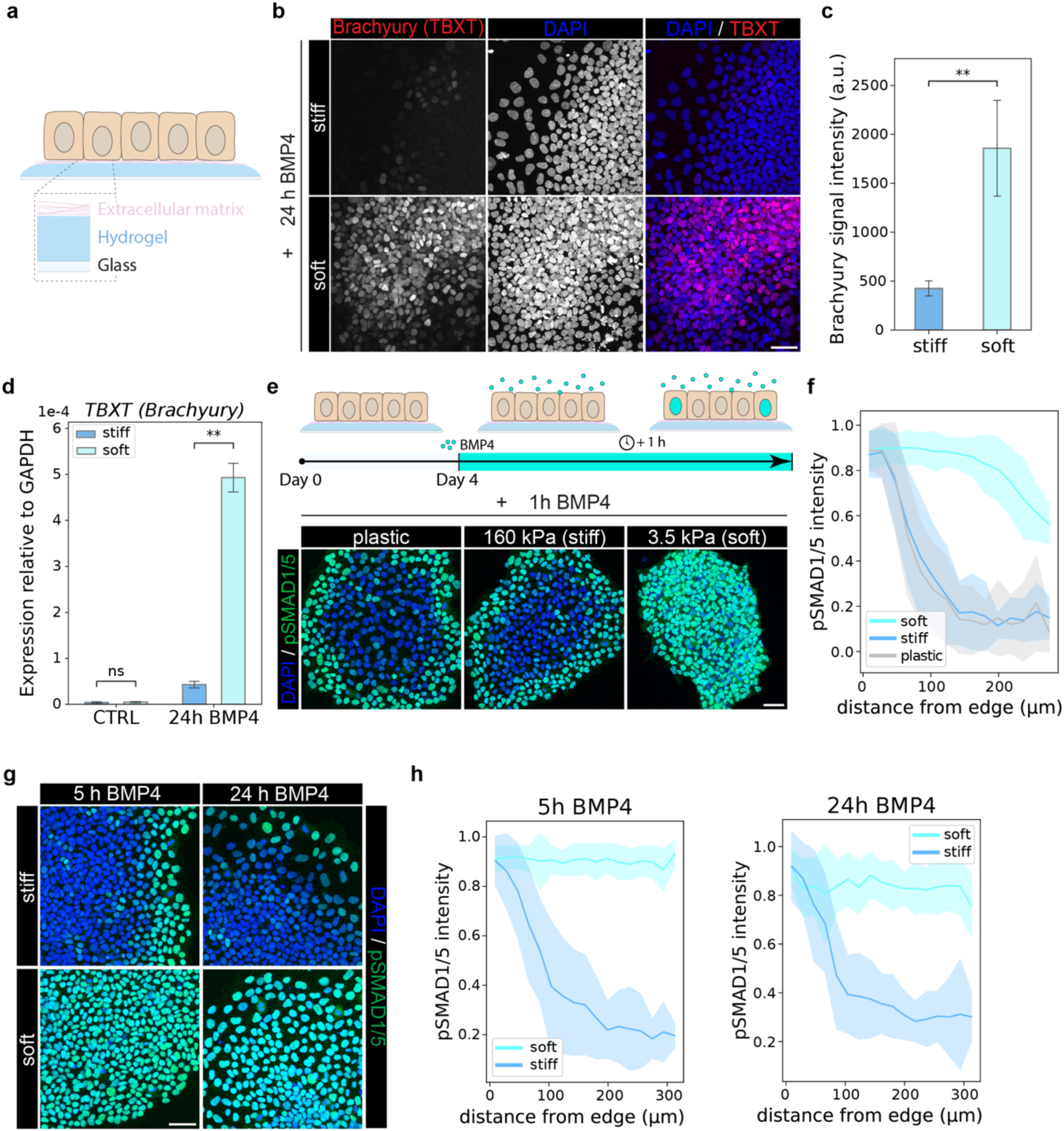
hPSCs on soft substrates lose spatially patterned response to BMP4. a. A schematic of the experimental system; hPSCs are cultured on hydrogels of various stiffnesses coated with vitronectin. b. Representative immunofluorescence images (maximum projection) of hPSCs on stiff and soft substrates stained for Brachyury (T) and DAPI after 24 h of BMP4 treatment. Scale bar = 50 μm. c. Quantification of Brachyury signal intensity across maximum projection images. P-value determined by Welch’s t-test. ** indicates p < 0.01. Here and everywhere in text and figure legends, N = 3 indicates individual data from three independent experiments representing biological replicates. d. qPCR data quantifying the level of expression of *TBXT* gene (*Brachyury*) normalised to the expression level of GAPDH in hPSCs on stiff and soft substrates, both with and without 24 h of BMP4 treatment. Individual data came from N = 3 independent experiments representing biological replicates. P-value determined by Welch’s t-test. ns indicates p > 0.05, ** indicates p < 0.01. e. Top: Schematic representing the experimental setup consisting of hPSCs cultured on stiff and soft substrates, exposed to 1 h of BMP4 treatment. Bottom: Representative immunofluorescence merged images (maximum projection) of hPSCs on plastic, stiff and soft substrates stained for pSMAD1/5 and DAPI after 1 h of BMP4 treatment. Scale bar = 50 μm. f. Quantification of pSMAD1/5 signal from immunofluorescence images of hPSCs on plastic, stiff and soft substrates after 1 h of BMP4 treatment. The signal intensity in each nucleus is plotted as a function of the distance of that nucleus from the colony edge, normalized by the 98^th^ percentile value of each image. The continuous line represents the mean signal intensity and edges of the shaded areas represent +/- standard deviation from N = 3 biological replicates. g. Representative immunofluorescence merged images (maximum projection) of hPSCs on stiff and soft substrates, treated with BMP4 for 5 h or 24 h and stained for pSMAD1/5 and DAPI. Scale bar = 50 μm. h. Quantification of pSMAD1/5 signal from immunofluorescence images of hPSCs on stiff and soft substrates after 5 h of BMP4 treatment and after 24 h of BMP4 treatment. The signal intensity in each nucleus is plotted as a function of the distance of that nucleus from the colony edge, normalized by the 98^th^ percentile value of each image. The continuous line represents the mean signal intensity and edges of the shaded areas represent +/- standard deviation from N = 3 biological replicates.

To address whether substrate stiffness affects the immediate response of hPSCs to BMP4, we investigated the phosphorylation level of the primary effector of the BMP4 signalling pathway, SMAD1/5, which is directly phosphorylated by the activated BMP receptor, BMPRI^34^. pSMAD1/5 levels were monitored in response to 1 h of BMP4 exposure, after 4 days of culture on soft and stiff hydrogels, as well as on plastic (Fig. 1e – schematic). When cultured on plastic, consistent with previous observations^6,19,35^, hPSCs responded to the BMP4 signal mainly in an outer ring of cells at the colony edge, spanning ∼40-50 μm from the edge (Fig. 1e,f). The same spatial pattern and level of response were observed when hPSCs were cultured on stiff hydrogels (Fig. 1e,f). However, when the cells were cultured on soft hydrogels, response to BMP4 signalling was observed across the entire colony (Fig. 1e,f). The loss of spatial restriction of the BMP4 response within colonies on soft hydrogels was preserved regardless of the colony size (Extended Data Fig. 1d). We also asked whether the difference in spatial response to BMP4 changed with longer exposure. The differential pattern of response between stiff and soft substrates remained unchanged both after 5 and 24 h of BMP4 treatment (Fig. 1g,h), leading to increased overall levels of pSMAD1/5 on soft substrates (Extended Data Fig. 1e).

Together, this data suggests that the mechanical cue generated by substrate stiffness strongly modulates patterned response of hPSCs to a biochemical signal, BMP4. On soft substrates, spatially restricted edge response is lost, which subsequently underpins increased mesodermal fate induction.

### Mechanosensing through FAK-PI3K activity regulates the patterning of BMP response

We next investigated the mechanism through which a change in substrate stiffness affects the patterning of BMP4 response in hPSC colonies. To determine if substrate stiffness primes hPSCs for differential response to BMP4, we first performed bulk RNA sequencing. Surprisingly, hPSCs cultured on stiff and soft substrates were undistinguishable at the gene expression level (Fig. 2a). In fact, only 62 genes were significantly differentially expressed between the two mechanical conditions (Extended Data Fig. 2a), and gene ontology term analysis did not highlight any specific gene categories related to response to developmental signals significantly enriched in either condition (Extended Data Fig. 2b). This analysis suggests that the change in response to BMP4 is not transcriptionally primed.

**Figure 2.**
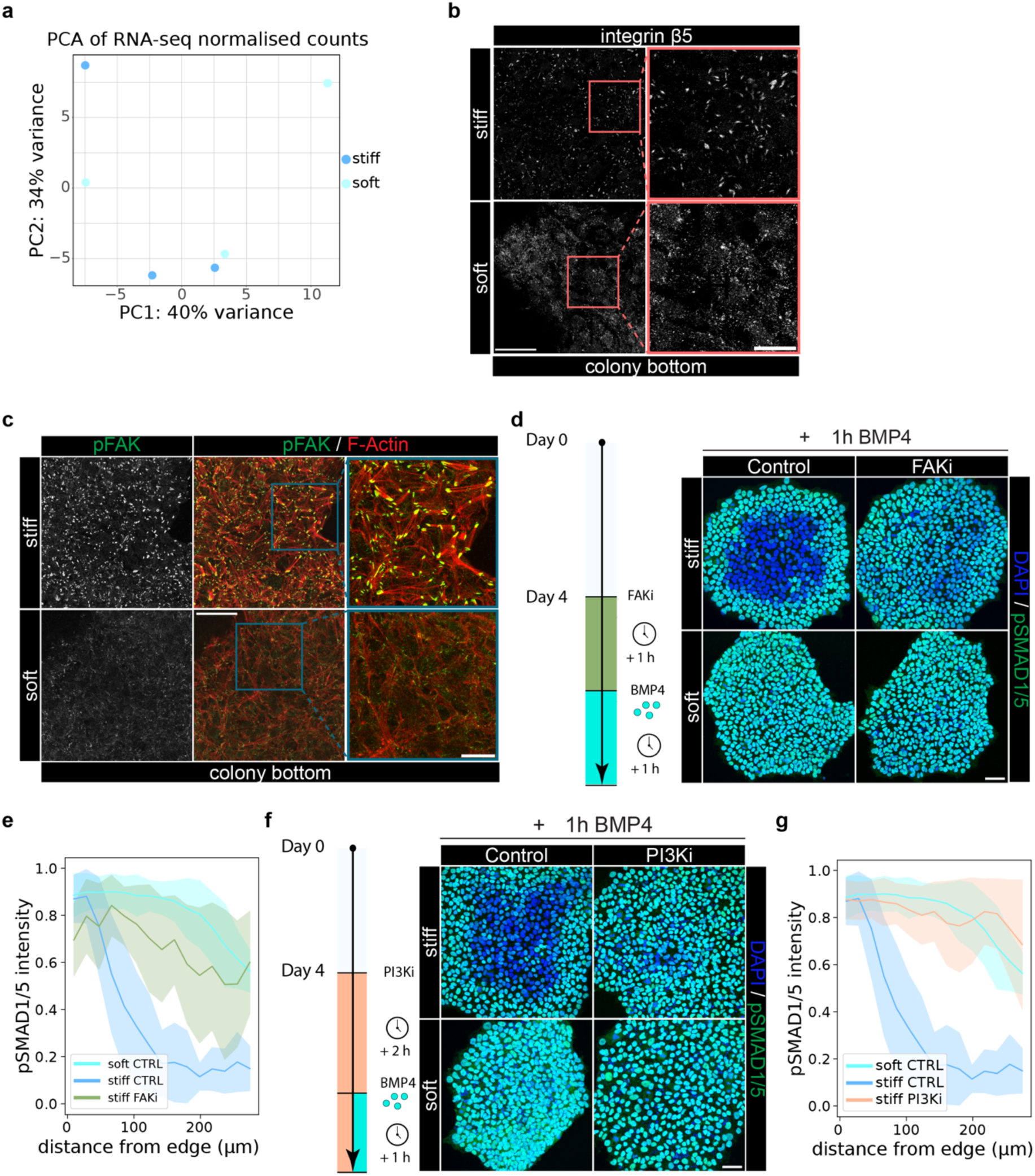
Reduced FAK-PI3K activity on soft substrates causes loss of patterning in BMP response. a. Principal component analysis of bulk RNA sequencing data from hPSCs on stiff and soft substrates after 4 days of culture, showing three biological replicates projected onto the first two principal components. b. Representative immunofluorescence images (maximum projection) of the colony bottom (bottom 5 confocal planes) of hPSCs on stiff and soft substrates, stained for integrin β5. Scale bar = 50 μm; zoomed in images scale bar = 20 μm. Individual data came from N = 3 independent experiments representing biological replicates. c. Representative immunofluorescence images (maximum projection) of colony bottom (bottom 5 confocal planes) of hPSCs on stiff and soft substrates, stained for pFAK and F-Actin. Scale bar = 50 μm; zoomed in images scale bar = 20 μm. Individual data came from N = 3 independent experiments representing biological replicates. d. Left: Schematic representing the experimental protocol involving FAK inhibition (FAKi) before the BMP4 treatment of hPSCs on stiff and soft substrates. Right: Representative immunofluorescence merged images (maximum projection) of hPSCs on stiff and soft substrates stained for pSMAD1/5 and DAPI after 1 h of BMP4 treatment, with or without the FAKi treatment. Scale bar = 50 μm. e. Quantification of pSMAD1/5 signal from immunofluorescence images of hPSCs in stiff control, soft control and stiff FAK inhibitor (FAKi) condition, after 1 h of BMP4 treatment. The signal intensity in each nucleus is plotted as a function of the distance of that nucleus from the colony edge, normalized by the 98^th^ percentile value of each image. The continuous line represents the mean signal intensity and edges of the shaded areas represent +/- standard deviation from N = 3 biological replicates. f. Left: Schematic representing the experimental protocol involving PI3K inhibition (PI3Ki) before the BMP4 treatment of hPSCs on stiff and soft substrates. Right: Representative immunofluorescence merged images (maximum projection) of hPSCs on stiff and soft substrates stained for pSMAD1/5 and DAPI after 1 h of BMP4 treatment, with or without the PI3Ki treatment. Scale bar = 50 μm. g. Quantification of pSMAD1/5 signal from immunofluorescence images of hPSCs in stiff control, soft control and stiff PI3K inhibitor (PI3Ki) condition, after 1 h of BMP4 treatment. The signal intensity in each nucleus is plotted as a function of the distance of that nucleus from the colony edge, normalized by the 98^th^ percentile value of each image. The continuous line represents the mean signal intensity and edges of the shaded areas represent +/- standard deviation from N = 3 biological replicates.

Cells primarily sense substrate stiffness through integrins^36^. Thus, to directly assess how substrate stiffness modulates the mechanosensing machinery in hPSCs, we compared transcription levels of genes encoding integrin subunits between stiff and soft substrates. We found that integrin genes were not differentially expressed between the two substrates (Extended Data Fig. 2c). However, we did observe that on the basal side of the cells, one of the most highly expressed integrin subunits, integrin β5, formed larger focal points on stiff substrates than on soft substrates (Fig. 2b). Clustering of integrins is a hallmark of adhesion maturation and generation of focal adhesions^37^, which causes activation of downstream intracellular effectors that regulate mechanosensing/ mechanotransduction, such as Focal Adhesion Kinase (FAK)^38,39^. Thus, we assessed FAK activity by investigating phosphorylation of FAK at Tyr 397^40^. We found that phosphorylated FAK (pFAK) displayed large clusters on the basal side of hPSC colonies on stiff substrates, while overall levels and cluster size of pFAK were reduced on soft substrates (Fig. 2c and Extended Data Fig. 2d). We further observed that F-actin formed strong stress fibres on stiff substrates, indicative of increased mechanotransduction, while stress fibres were substantially reduced on soft substrates (Fig. 2c). Together, these data unsurprisingly indicate decreased mechanosensing and mechanotransduction on soft substrates.

We then asked whether FAK activity played a role in BMP4 response differences observed between soft and stiff substrates. To investigate this, we restricted FAK activity in hPSCs using the FAK Inhibitor 14, which inhibits phosphorylation at Tyr 397^41^ (Fig. 2d - schematic). We found that, upon FAK inhibition, the response to BMP4 on stiff substrates occurred throughout the colony, phenotypically matching the response normally seen on soft substrates (Fig. 2d,e). We checked using RNA sequencing that this effect, on this timescale, was not due to changes in expression of genes involved in pathways relevant for BMP4 response (Extended Data Fig. 2e). One of the most well-described targets of FAK is PI3K^42^. We thus asked whether the PI3K activity is involved in the differences in response to BMP4 observed between soft and stiff substrates. Inhibition of PI3K activity with LY294002^43^ also caused loss of the spatial patterning of BMP4 response on stiff substrates in comparison to the non-treated stiff condition (Fig. 2f,g), phenotypically matching the response we observed on soft substrates (Fig. 2f,g). Again, RNA sequencing data indicated that PI3K inhibition on stiff substrates, on this timescale, did not cause a significant change in expression of genes controlling the pathways relevant for BMP4 response (Extended Data Fig. 2f). Finally, we asked if increasing PI3K activity would conversely restrict the spatially patterned BMP4 response, and inhibited a negative regulator of PI3K, PTEN ^44,45^. We found that PTEN inhibition further restricted the spatial response to BMP4 on stiff substrates and led to generation of a spatially patterned response on soft substrates (Extended Data Fig. 2g,h).

Together, this data suggests that the loss of spatial restriction of BMP4 response in hPSCs on soft substrates is not driven by changes in gene expression levels but is the result of reduced mechanical signalling through the FAK-PI3K pathway. Since the FAK-PI3K cascade is known to be implicated in regulating epithelial structure and polarity^46^, and given that BMPRs display polarised epithelial localisation^6^, we next asked whether substrate stiffness influences hPSC epithelial organisation.

### Increased epithelial permeability in hPSCs on soft substrates causes the loss of patterning in BMP response

To investigate whether changes in epithelial organisation contributed to the loss of patterned response to BMP4 on soft substrates, we first compared epithelial apical geometry in hPSCs on soft and stiff substrates. We stained cells for the tight junction protein ZO-1, which outlines apical domains. With ZO-1 staining, we observed more heterogeneous and irregularly shaped apical domains on soft compared to stiff substrates (Fig. 3a). We then quantified the apical cell shape index (CSI), which quantifies deviation from a circular shape, and found that the apical CSI was significantly higher for cells on soft compared to stiff substrates (Fig. 3b), confirming that hPSCs displayed more elongated and irregular apical shapes on soft substrates. PI3K inhibition on stiff substrates also led to more irregularly shaped apical domains (Fig. 3a), and the apical CSI increased to levels comparable to hPSCs on soft substrates (Fig. 3b).

**Figure 3.**
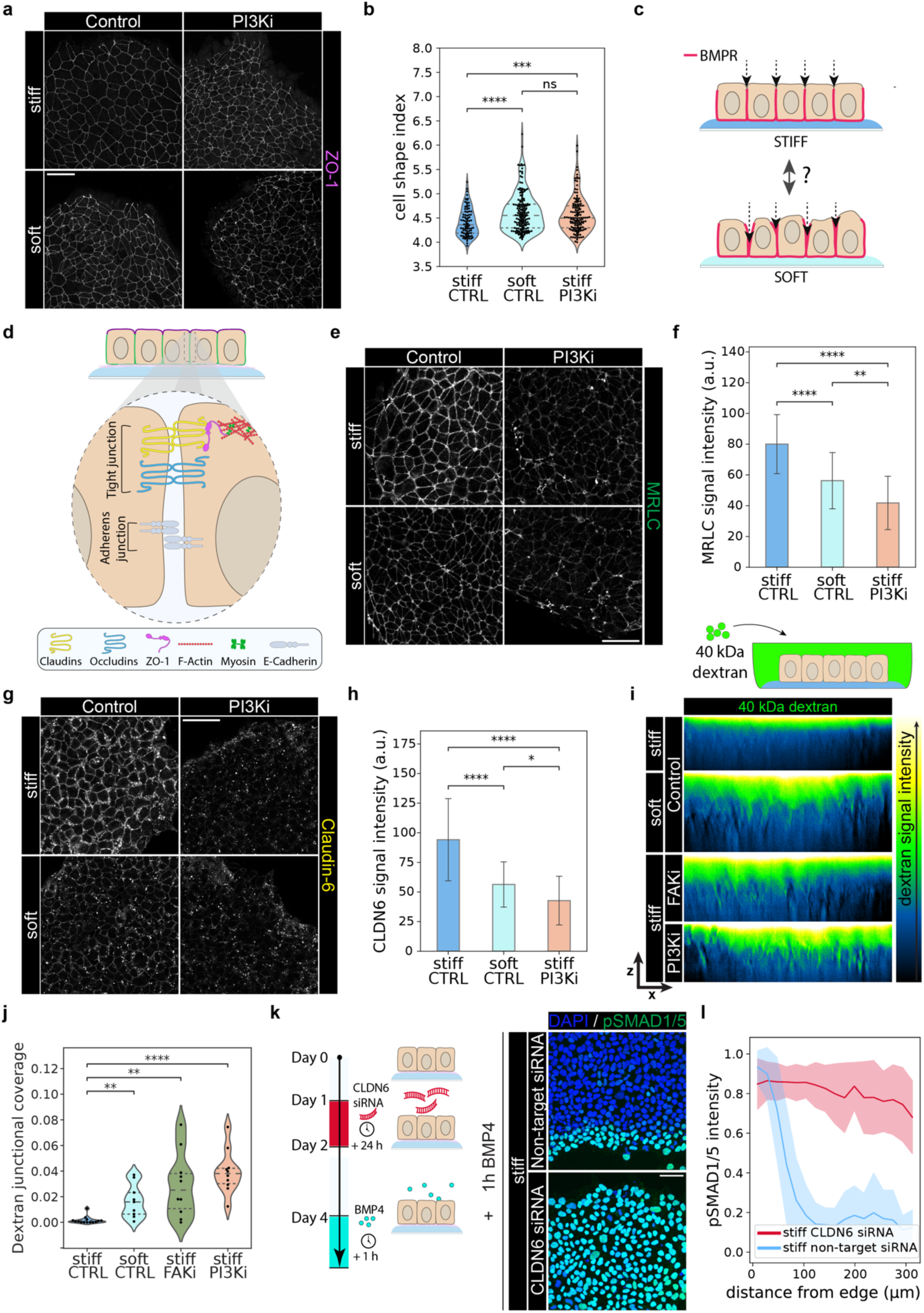
Loss of junctional integrity on soft substrates abrogates spatially restricted response to BMP. a. Representative immunofluorescence images (maximum projection) of colony top (top 5 confocal planes) of hPSCs on stiff and soft substrates, with or without PI3K inhibitor (PI3Ki) treatment, stained for ZO-1. Scale bar = 50 μm. b. Violin plots of the cell shape index (CSI) of apical surfaces of the cells in stiff control, soft control and stiff PI3K inhibitor (PI3Ki) conditions. 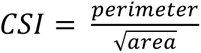. CSI is lowest for the circle (≈ 3.54), and any shape that deviates from a circle will have a higher CSI. Each point represents a single cell. Individual data came from N = 3 independent experiments representing biological replicates. P-value determined by Welch’s t-test. ns indicates p > 0.05, *** indicates p < 0.001, and **** indicates p < 0.0001. c. Schematic illustrating the hypothesis that substrate stiffness affects junctional integrity, which could in turn affect accessibility of the BMPR to the BMP4 ligand. d. Schematic showing components of tight and adherens junctions in an epithelium and the division of the epithelial cell membranes into two domains: apical (purple) and basolateral (green). e. Representative immunofluorescence images (maximum projection) of colony top (top 5 confocal planes) of hPSCs on stiff and soft substrates, with or without PI3K inhibitor (PI3Ki) treatment, stained for MRLC. Scale bar = 50 μm. f. Quantification of MRLC signal intensity across maximum projection images. Individual data came from N = 3 independent experiments representing biological replicates. P-value determined by Welch’s t-test. ** indicates p-value < 0.01, **** indicates p-value < 0.0001. g. Representative immunofluorescence images (maximum projection) of colony top (top 5 confocal planes) of hPSCs on stiff and soft substrates, with or without FAK inhibitor (FAKi) or PI3K inhibitor (PI3Ki) treatment, stained for Claudin-6. Scale bar = 50 μm. h. Quantification of Claudin-6 signal intensity across maximum projection images. Individual data came from N = 3 independent experiments representing biological replicates. P-value determined by Welch’s t-test. * indicates p-value < 0.05, **** indicates p-value < 0.0001. i. Top: Schematic illustrating the permeability experiment using fluorescently labelled dextran of the size of the BMP4 ligand, designed to test the hypothesis of junctional integrity change with substrate stiffness. Bottom: Representative images of fluorescent dextran signal along the apicobasal axis (x-z plane) on stiff and soft substrates, in the control condition or after the FAK inhibitor (FAKi) or PI3K inhibitor (PI3Ki) treatment. The lookup table indicates dextran signal intensity from low (blue) to high (white). j. Quantification of dextran signal at the lateral junctions, defined as the fraction of the central 50% of the lateral junction mask (green in Extended Data Fig. 3c) covered by the thresholded (brightest) signal pixels (red in Extended Data Fig. 3c). See *Materials and Methods* for further details about quantification. Individual data came from N = 3 independent experiments representing biological replicates. P-value determined by Welch’s t-test. ** indicates p-value < 0.01, **** indicates p-value < 0.0001. k. Left: Schematic representing the experimental protocol involving Claudin-6 knockdown using siRNAs in hPSCs on stiff substrates before the BMP4 treatment. Right: Representative immunofluorescence merged images (maximum projection) of hPSCs on stiff substrates treated with Claudin-6 targeting or non-targeting siRNAs, stained for pSMAD1/5 and DAPI after 1 h of BMP4 treatment. Scale bar = 50 μm. l. Quantification of pSMAD1/5 signal from immunofluorescence images of hPSCs on stiff substrates, treated with non-targeting siRNAs or Claudin-6 targeting siRNAs, after 1 h of BMP4 treatment. The signal intensity in each nucleus is plotted as a function of the distance of that nucleus from the colony edge, normalized by the 98^th^ percentile value of each image. The continuous line represents the mean signal intensity and edges of the shaded areas represent +/- standard deviation from N = 3 biological replicates.

We then asked whether the observed differences in epithelial morphology between soft and stiff substrates contribute to the differential response to BMP4 on the two types of substrates. BMPRs in hPSCs have been previously shown to be localised on the basolateral membrane of cells^6,47^ and epithelial junctions restrict accessibility of the basolateral membrane to soluble molecules^48^. This decreased basolateral accessibility limits the response to apically applied BMP4 in colony centre^6^. Since we observed loss of spatial restriction of response to BMP4 on soft substrates, we hypothesised that integrity of epithelial junctions on soft substrates was reduced (Fig. 3c).

Epithelial integrity is maintained by tight junctions which are primarily composed of claudins and occludins that generate a barrier to soluble particles between the cells, and ZO proteins that connect the junction to the actomyosin cytoskeleton^48^ (Fig. 3d). Myosin exerts tension on tight junctions^49^, and changes in tension can result in displacement of junctional proteins and loss of integrity^50–52^. We thus visualised junctional Myosin Regulatory Light Chain (MRLC) and observed a significant reduction of junctional MRLC on soft compared to stiff substrates (Fig. 3e,f). Moreover, we found that PI3K inhibition strongly reduced junctional MRLC levels on stiff substrates (Fig. 3e,f). Finally, inhibition of myosin activity using blebbistatin caused a loss in spatially patterned response to BMP4 on stiff substrates (Extended Data Fig. 3a), suggesting that reduction of junctional tension contributed to enhanced BMP4 response.

Reduction in junctional myosin on soft substrates prompted us to investigate the effect of substrate stiffness on junctional proteins. We observed no clear difference in ZO-1 levels between substrates (Fig. 3a), so we investigated another group of essential tight junction proteins, claudins^53^. hPSCs highly express one form of claudin, *CLDNC*^54^ (Extended Data Fig. 3b). Therefore, we compared Claudin-6 expression between substrates of different stiffnesses. We found no significant difference in gene expression levels (Extended Data Fig. 3b) but observed a significant reduction in junctional levels of Claudin-6 on soft substrates in comparison to stiff substrates (Fig. 3g,h). PI3K inhibition on stiff substrates led to a reduction in junctional Claudin-6 down to levels of Claudin-6 comparable to soft substrates (Fig. 3g,h). Together, these observations suggest that tight junctions are under lower contractile tension and accumulate lower levels of junctional proteins on soft compared to stiff substrates, suggesting that junctional integrity might be reduced on soft substrates.

To directly assess junctional integrity and, as a result, junctional permeability of hPSC epithelia, we apically delivered fluorescently labelled dextran of a size roughly matching the size of the BMP4 ligand (40 kDa) (Fig. 3i - schematic). We then performed live imaging to assess the extent to which dextran penetrates the hPSC epithelia on the different substrates (Extended Data Fig. 3c,d; see *Materials and Methods*). We found that hPSCs cultured on soft substrates exhibited a significantly higher penetration of dextran into the epithelial junctions (Fig. 3i,j). Similarly enhanced dextran penetration within the epithelium was observed when cells on stiff substrates were treated with the FAK inhibitor or the PI3K inhibitor prior to dextran addition (Fig. 3i,j). Altogether, our data show that the hPSC epithelium on soft substrates exhibits higher permeability to BMP4-sized molecules, indicating lower junctional integrity.

We next asked whether decreasing junctional integrity in hPSCs on stiff substrates is sufficient to abrogate the spatially restricted response to BMP4. To test this, we knocked down Claudin-6 in hPSCs on stiff substrates using siRNAs (Extended Data Fig. 3e), and monitored their response to BMP4 (Fig. 3k - schematic). We found that hPSCs depleted for Claudin-6 responded to BMP4 throughout the colony on stiff substrates, phenocopying the response observed on soft substrates (Fig. 3k,l).

Together, these data suggest that on soft substrates, hPSCs exhibit reduced junctional integrity compared to cells on stiff substrates, which is sufficient to lose spatially regulated response to BMP4.

### Disrupted epithelial polarity in hPSCs on soft substrates contributes to the loss of patterning in BMP response

Given the received knowledge that BMPRs are basolaterally localised, loss of junctional integrity on soft substrates could explain loss of spatially regulated response to BMP4. However, loss of the spatially restricted pattern of BMP4 response could also be caused by redistribution of receptors to the apical side, which would then be directly exposed to the signal. To investigate this alternate possibility, we assessed the localisation of BMPRs in both conditions. To do this, we first compared the localisation of BMPR1A, the main BMPR1 paralog expressed in hPSCs (Extended Data Fig. 4a), between stiff and soft substrates. We found that BMPR1A localised predominantly to the basal and lateral sides of the hPSCs on stiff substrates but displayed a strongly increased apical distribution in cells cultured on soft substrates (Fig. 4a-c; see *Materials and Methods*). This suggests that on soft substrates, BMPR1A partially re-localises to the apical membrane domain.

**Figure 4.**
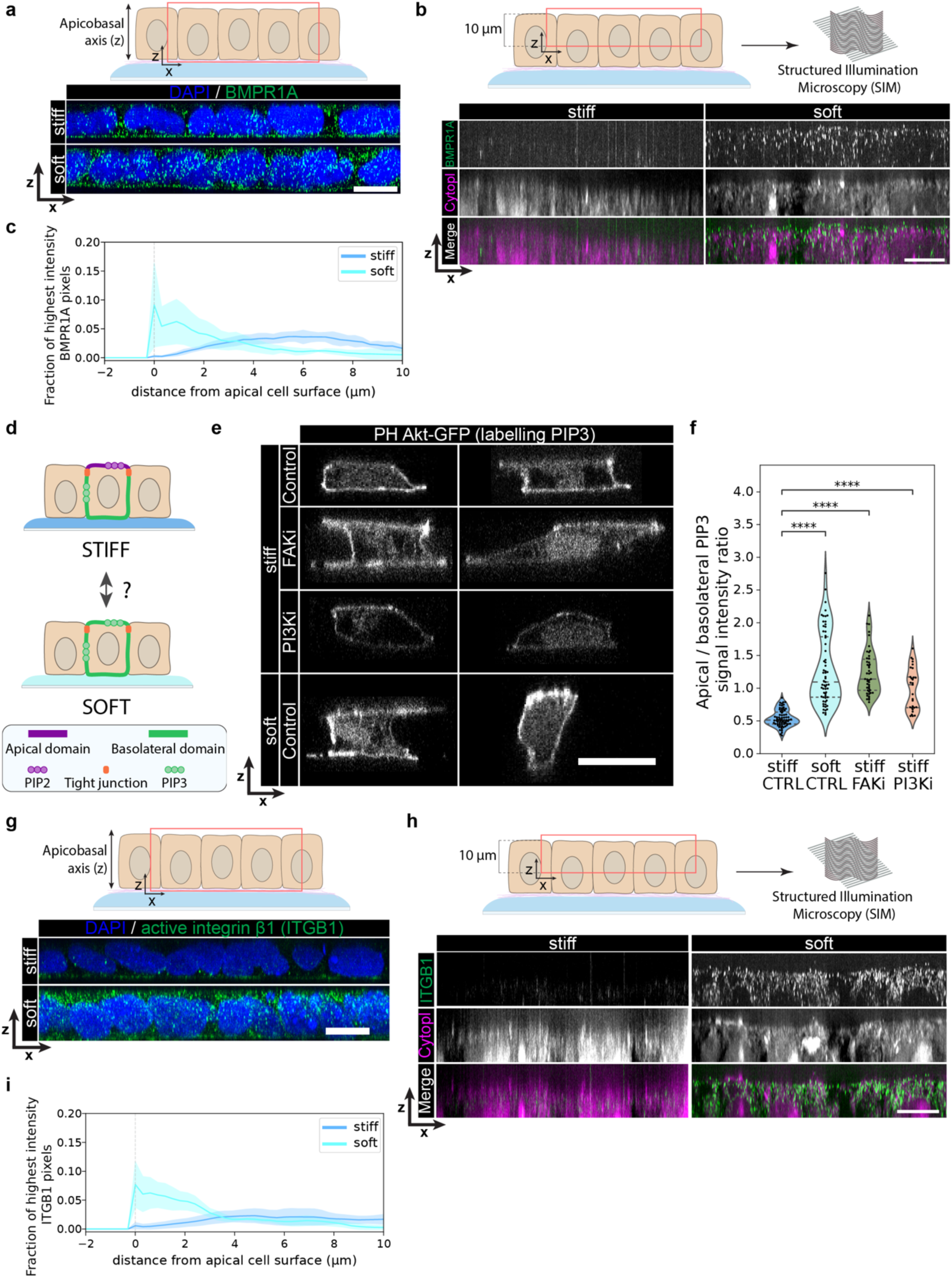
Disrupted polarity in hPSCs on soft substrates causes the loss of patterned response to BMP4. a. Top: Schematic showing the part of the hPSC colonies that was imaged using confocal microscopy (red box). Bottom: Representative immunofluorescence merged images (maximum projection) of hPSCs on stiff and soft substrates, stained for BMPR1A and DAPI. Images were reconstructed in x-z plane (along the apicobasal axis); 50 imaging planes were projected in y-axis. Scale bar = 15 μm. b. Top: Schematic showing the part of the hPSC colonies that was imaged using structured illumination microscopy (SIM). Bottom: Representative immunofluorescence images (maximum projection) of hPSCs on stiff and soft substrates, stained for BMPR1A and with CellMask HCS dye staining the cytoplasm. Images were reconstructed in x-z (i.e., along the apicobasal axis), and the top 10 μm of the apicobasal axis were imaged; 40 imaging planes were projected in y-axis. Scale bar = 10 μm. c. Plot of the fraction of BMPR1A-positive pixels against the distance from cell surface from SIM images. The continuous line represents the mean signal intensity and edges of the shaded areas represent +/- standard deviation from N = 3 biological replicates. d. Schematic of apical and basolateral membrane domains in epithelial hPSCs, defined by the localisation of phospholipids PIP2 and PIP3. Schematic also represents the hypothesis of polarity disruption as a consequence of substrate stiffness change. e. Representative images of hPSCs on stiff and soft substrates in control conditions and upon FAK inhibitor (FAKi) or PI3K inhibitor (PI3Ki) treatment, transfected with the plasmid encoding the GFP-tagged PH domain of Akt. Images were reconstructed in the x-z plane, i.e., along the apicobasal axis. Scale bar = 20 μm. f. Quantification of the apical/basolateral ratio of PH Akt-GFP (labelling PIP3) signal intensity in individual hPSCs on stiff and soft substrates in control condition as well as on stiff substrates upon FAK inhibitor (FAKi) or PI3K inhibitor (PI3Ki) treatment. See Extended Data Fig. 4b for quantification pipeline. Individual data came from N = 3 independent experiments representing biological replicates. P-value determined by Welch’s t-test. **** indicates p-value < 0.0001. g. Top: Schematic showing the part of the hPSC colonies that was imaged using confocal microscopy (red box). Bottom: Representative immunofluorescence merged images (maximum projection) of hPSCs on stiff and soft substrates, stained for the active form of integrin β1 and DAPI. Images were reconstructed in x-z plane (along the apicobasal axis); 50 imaging planes were projected in y-axis. Scale bar = 15 μm. h. Top: Schematic showing the part of the hPSC colonies that was imaged using structured illumination microscopy (SIM) (red box). Bottom: Representative immunofluorescence images (maximum projection) of hPSCs on stiff and soft substrates, stained for the active form of integrin β1 and with HCS dye staining the cytoplasm. Images were reconstructed in x-z (i.e., along the apicobasal axis), and the top 10 μm of the apicobasal axis were imaged; 20 imaging planes were projected in the y-axis. Scale bar = 10 μm. i. Plot of the fraction of active integrin β1-positive pixels against the distance from the cell surface from SIM images. The continuous line represents the mean signal intensity and edges of the shaded areas represent +/- standard deviation from N = 3 biological replicates.

Distribution of membrane proteins to apical and basolateral membrane domains is controlled by epithelial apical-basal polarity, maintained by distinct polarity proteins and lipids in each domain (Fig. 4d)^55^. Since we observed localisation of conventionally basolateral BMPR1A at the apical membrane on soft substrates, we asked whether cells on soft substrates displayed disrupted epithelial polarity (Fig. 4d). To address this question, we investigated localisation of phosphatidylinositol (3,4,5)-trisphosphate (PIP3), a phospholipid which in polarised epithelia predominantly localises to the basolateral membrane domain^56,57^ (Extended Data Fig. 4b). While on stiff substrates PIP3 indeed predominantly localised basolaterally (Fig. 4e,f), on soft substrates, hPSCs exhibited a strong increase in PIP3 at the apical membrane (Fig. 4e,f). Likewise, PIP3 also displayed enhanced apical localisation in hPSCs on stiff substrates treated with FAK inhibitor and PI3K inhibitor (Fig. 4e,f). Together, these observations suggest that reduced mechanosensing/mechanotransduction in hPSC on soft substrates leads to a general disruption of apical membrane identity.

Apicobasal axis orientation and polarisation are initiated at least in part by activation of integrins in response to binding to the ECM, specifying the basal side of the cells^58^. We thus asked whether active integrin localisation differed between stiff and soft substrates. The most highly expressed integrin subunit in hPSCs, integrin β1 (Extended Data Fig. 2c), was active predominantly on the basal side of the cells on stiff substrates (Fig. 4g). However, on soft substrates we observed increased localisation of active integrin β1 to the apical surface, while not being detectable on the apical membrane on stiff substrates (Fig. 4g-i).

These data suggest that hPSCs on soft substrates exhibit disrupted apico-basal polarity, which could explain increased localisation of the BMPR to the apical domain. Taken altogether, our observations suggest a multifaceted mechanism: decreased junctional integrity combined with disrupted epithelial polarity increases BMPR exposure throughout the epithelium, which in turn leads to a loss of spatial restriction of BMP4 response in hPSCs on soft substrates.

### Apical laminin retention causes epithelial reorganisation and loss of spatial patterning in BMP response

The fact that hPSCs on soft substrates exhibit apically active integrin β1 prompted us to investigate the potential presence of an ECM protein at the apical cell surface, which could cause apical activation of integrin β1. hPSCs are known to secrete laminin^59^, which is, alongside collagen IV, a primary constituent of the basement membrane to which hPSC-equivalent epiblast cells are attached to *in vivo*^32^. Hence, we stained for laminin in our different culture conditions and observed an increased presence of laminin on the apical side of hPSCs on soft substrates in comparison to stiff substrates (Fig. 5a,b). Yet, no laminin subunit was differentially expressed in hPSCs between the stiff and soft substrates (Extended Data Fig. 5a). This suggests that hPSCs on soft substrates might display enhanced apical retention of secreted laminin compared to cells cultured on stiff substrates.

**Figure 5.**
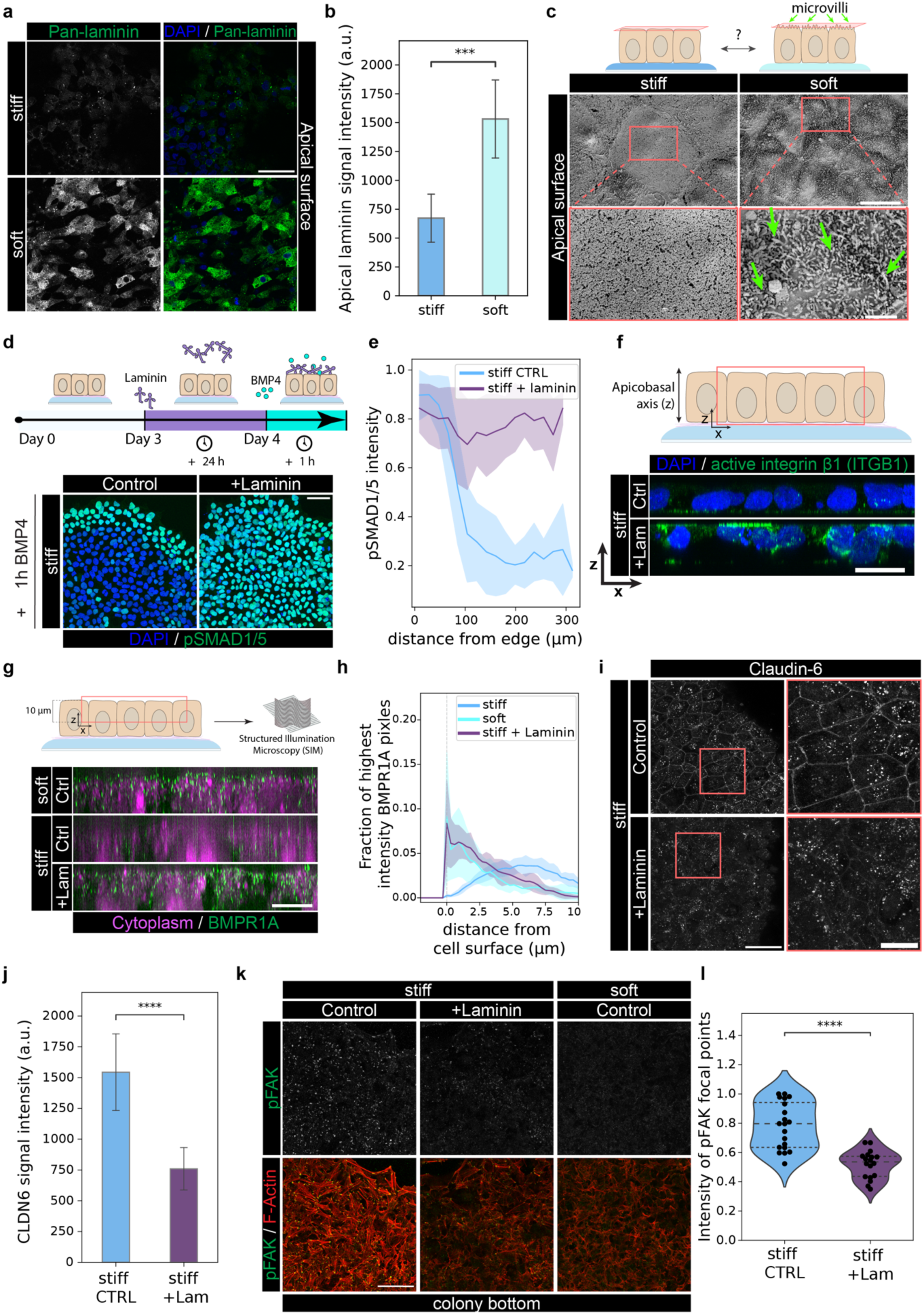
Apical laminin accumulation disrupts epithelial polarity and causes loss of spatially patterned response to BMP4. a. Representative immunofluorescence images of the colony apical surface of hPSCs on stiff and soft substrates, without permeabilization, stained for pan-laminin and DAPI. Scale bar = 50 μm. b. Quantification of apical laminin signal intensity across images. Individual data came from N = 3 independent experiments representing biological replicates. P-value determined by Welch’s t-test. *** indicates p-value < 0.001. c. Top: Schematic of the hypothesized apical cell surface in hPSCs on stiff and soft substrates. Green arrows point to microvilli, red rectangle represents the plane imaged with SEM. Bottom: SEM images of the apical surface of hPSCs on stiff and soft substrates. Green arrows point to individual microvilli. Scale bar = 10 μm; zoomed in images scale bar = 2 μm. Representative images came from N = 3 independent experiments representing biological replicates. d. Top: Schematic representing the experimental protocol involving soluble laminin addition into the cell medium of hPSCs on stiff substrates before the BMP4 treatment. Bottom: Representative immunofluorescence merged images (maximum projection) of hPSCs on stiff substrates stained for pSMAD1/5 and DAPI after 1 h of BMP4 treatment, with or without addition of soluble laminin into the cell medium. Scale bar = 50 μm. e. Quantification of pSMAD1/5 signal from immunofluorescence images of hPSCs on stiff substrates, with or without the addition of soluble laminin into the cell medium and after 1 h of BMP4 treatment. The signal intensity in each nucleus is plotted as a function of the distance of that nucleus from the colony edge, normalized by the 98^th^ percentile value of each image. The continuous line represents the mean signal intensity and edges of the shaded areas represent +/- standard deviation from N = 3 biological replicates. f. Top: Schematic showing the part of the hPSC colonies that was imaged using confocal microscopy (red box). Bottom: Representative immunofluorescence merged images (maximum projection) of hPSCs on stiff substrates, with or without the addition of soluble laminin, stained for the active form of integrin β1 and DAPI. Images were reconstructed in the x-z plane (along the apicobasal axis); 50 imaging planes were projected in y-axis. Scale bar = 20 μm. g. Top: Schematic showing the part of the hPSC colonies that was imaged using structured illumination microscopy (SIM). Bottom Representative immunofluorescence merged images (maximum projection) of hPSCs on stiff and soft substrates, with or without the addition of soluble laminin, and stained for BMPR1A and with HCS dye staining the cytoplasm. Images were reconstructed x-z (i.e., along the apicobasal axis), and the top 10 μm of the apicobasal axis was imaged; 40 imaging planes were projected in y-axis. Scale bar = 10 μm. h. Plot of the fraction of BMPR1A-positive pixels against the distance from the cell surface. The continuous line represents the mean signal intensity and edges of the shaded areas represent +/- standard deviation from N = 3 biological replicates. i. Representative immunofluorescence images (maximum projection) of colony top (top 5 confocal planes) of hPSCs on stiff substrates, with or without soluble laminin addition into the medium, stained for Claudin-6. Scale bar = 50 μm; zoomed in images scale bar = 20 μm. j. Quantification of junctional Claudin-6 signal intensity across maximum projection images. Individual data came from N = 3 independent experiments representing biological replicates. P-value determined by Welch’s t-test. **** indicates p-value < 0.0001. k. Representative immunofluorescence images (maximum projection) of colony bottom (bottom 5 confocal planes) of hPSCs on stiff and soft substrates, in control condition or upon addition of soluble laminin into the cell medium, stained for pFAK and F-Actin. Scale bar = 50 μm. l. Quantification of normalised signal intensity in pFAK focal points in hPSCs on stiff substrates in control condition and upon addition of soluble laminin into the cell medium. Individual data came from N = 3 independent experiments representing biological replicates. P-value determined by Welch’s t-test. **** indicates p-value < 0.0001.

To investigate what could lead to increased laminin apical retention on soft substrates, we assessed the topology of the apical surface using scanning electron microscopy. Notably, hPSCs on soft substrates exhibited a very high density of microvilli on their apical surfaces compared to stiff substrates, where microvilli were sparse or absent altogether (Fig. 5c). Furthermore, in cells on soft substrates, we also observed increased levels of phosphorylated Ezrin-Radixin-Moesin (p-ERM) proteins (Extended Data Fig. 5b,c), which are known to be enriched in microvilli^60^. Thus, the apical surface of hPSCs on soft substrates appears to display a higher density of membrane microvilli, which might contribute to the retention of apical ECM components^61,62^.

We then asked whether apical laminin presence was sufficient to abrogate spatially restricted response to BMP4. To test this hypothesis, we treated hPSCs cultured on stiff substrates with soluble laminin in the medium before exposing them to BMP4 (Fig. 5d - schematic). We checked that laminin added to the media is indeed retained on the apical surfaces of hPSCs on stiff substrates (Extended Data Fig. 5d). We then asked whether apical laminin exposure affected BMP4 response. Strikingly, we observed that hPSCs on stiff substrates exposed to apical laminin displayed high levels of pSMAD1/5 across the entire colony in response to BMP4 treatment (Fig. 5d,e), recapitulating the phenotype observed in hPSCs on soft substrates (Fig. 1e). Moreover, soluble laminin treatment of hPSCs on stiff substrates led to increased apical localisation of active integrin β1 (Fig. 5f). We thus asked whether integrin binding to apical laminin mediated the observed loss of spatially patterned response to BMP4. Integrin β1 heterodimerises to bind laminin and the most expressed binding partner is integrin α6 (Extended Data Fig. 2c). Therefore, we used blocking antibodies to prevent binding of integrin β1 and α6 to laminin prior to and during soluble laminin treatment and monitored the response to BMP4 (Extended Data Fig. 5e - schematic). We found that blocking integrin β1 reduced the response to BMP4 in the colony centre, and blocking integrin α6 and β1 together completely reversed the effects of apical laminin addition (Extended Data Fig. 5e,f).

Next, we investigated whether the enhanced response to BMP4 in hPSCs on stiff substrates following apical laminin exposure was mediated by the same mechanisms as those promoting BMP4 response on soft substrates. We found that apical exposure to laminin strongly increased localisation of BMPR1A to the apical side of hPSCs on stiff substrates, resulting in a BMPR distribution comparable to that observed in cells on soft substrates (Fig. 5g,h), suggesting disruption of epithelial polarity. Furthermore, we observed that laminin treatment of hPSCs on stiff substrates caused a significant reduction of junctional Claudin-6 localisation (Fig. 5i,j). These observations indicate that apical exposure to laminin in hPSCs on stiff substrates phenocopies the BMPR redistribution and reduced junctional integrity observed in hPSCs on soft substrates (Fig. 3 and 4).

Finally, given the observed phenocopying of the soft substrate, we asked whether apical laminin accumulation also perturbs basal mechanosensing of stiff substrates. To address this, we assessed FAK activity, whose mechanosensing activity correlates with and is dependent on substrate stiffness (Fig. 2c). Notably, apical laminin exposure in hPSCs on stiff substrates caused a reduction in basal FAK activity and a decrease in stress fibre formation (Fig. 5k,l). Together, this data suggests that apical ECM accumulation phenocopies soft substrates, showing consequential effects on epithelial organisation and BMP4 signalling, and possibly overriding mechanosensing.

### Softening of the basement membrane induces premature ectopic mesodermal differentiation in the mouse embryo

Finally, we asked whether substrate softening would also affect the patterning of BMP4 response *in vivo*. In the mouse embryo, epithelialized pluripotent epiblast cells are found at E5.0-5.5 of development (Fig. 6a – E5.5). Even though at E5.5, the apical surfaces of epiblast cells are exposed to the BMP4 signal present in the pro-amniotic cavity, they only begin to differentiate in response to BMP4 at E6.5 at the start of gastrulation, which causes epithelial-to-mesenchymal transition, and subsequent ingression through primitive streak (Fig. 6a – E6.5)^16^. The start of gastrulation has been shown to correlate with matrix metalloproteinase (MMP)-driven remodelling of the Collagen IV-rich basement membrane underlying the epiblast^31^. Therefore, we hypothesised that remodelling-induced softening of the basement membrane could potentiate the response of epiblast cells to BMP4 and mesodermal differentiation, like we observed in hPSCs on soft substrates *in vitro* (Fig. 1b-f). Basement stiffness is largely determined by collagen IV crosslinking^63–65^, controlled by lysyl oxidase^66^. We thus dissected E5.5 mouse embryos and treated them with a lysyl oxidase inhibitor, BAPN, to induce basement membrane softening (Fig. 6b). Embryos were treated for no longer than 16 h, so that the natural start of gastrulation at E6.5 (Fig. 6c - E6.5) is not yet reached. We then assessed mesodermal differentiation in the epiblast by staining for the early mesoderm lineage marker, Brachyury. Strikingly, while control embryos were mostly Brachyury negative at E5.5 + 16 hours (1 out of 15 embryos displayed Brachyury+ cells), 12 out of 27 embryos treated with BAPN exhibited Brachyury+ cells (Fig. 6c,d). Furthermore, Brachyury+ cells were present throughout the epiblast, not just restricted to the proximal-posterior epiblast as normally observed. These data demonstrate that basement membrane softening is sufficient to initiate premature Brachyury expression, and loss of spatial restriction in the pattern of fate acquisition. Together, this suggests that spatiotemporal response to developmental signals, which leads to fate patterning during mouse gastrulation, is conditioned by the basement membrane mechanics *in vivo*, consistent with our findings in the hPSCs *in vitro*.

**Figure 6.**
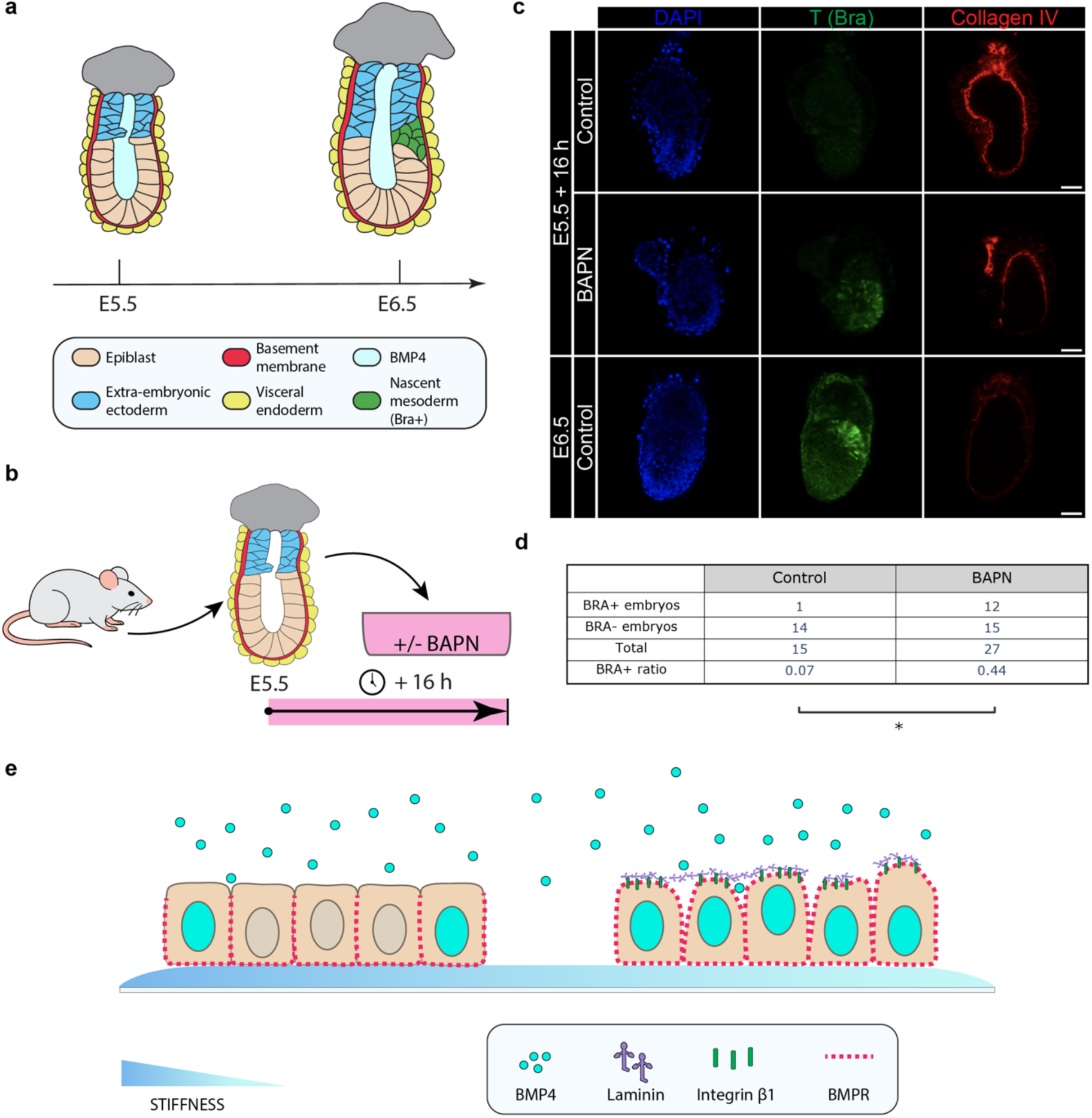
Softening the basement membrane induces premature and ectopic mesodermal differentiation in the mouse epiblast. a. Schematic depicting the anatomy of mouse embryos after implantation, at E5.5 and E6.5. b. Schematic showing the experimental setup involving treatment of dissected E5.5 mouse embryos with a LOX inhibitor, BAPN, that prevents Collagen IV crosslinking. c. Representative immunofluorescence images of E5.5 mouse embryos in control conditions and after 16 h of BAPN treatment, as well as control E6.5 embryos. Scale bar = 50 μm. d. Quantification of the fraction of embryos containing Brachyury+ cells at E5.5 in total number of E5.5 embryos. Individual data came from N = 3 independent experiments representing biological replicates. P-value = 0.0148, determined by Fisher’s exact test. e. Schematic depicting the mechanism of epithelial reorganization upon substrate softening leading to a change in spatial response to BMP4 in hPSCs.

## DISCUSSION

In this study, we provide mechanistic insight into how basement membrane mechanics regulates the spatial response to developmental signalling in hPSCs, highlighting the importance of understanding the crosstalk between mechanical and biochemical signalling. Mechanical and biochemical signalling have been mostly investigated in isolation, and their integration has been largely neglected in studies of cell fate choice, developmental patterning and morphogenesis. Our study demonstrates that changing basement membrane mechanics can regulate – either augment or limit – cellular response to instructive signals, unveiling a key and unappreciated mechanism for the control of developmental patterning.

We show that changing substrate stiffness changes the pattern of response of hPSCs to BMP4 signalling. On soft substrates, the BMP4 response is present across the whole colony as opposed to the marginal edge response observed on stiff substrates described previously^6,19,35^. The loss of a spatially restricted response eventually impacts individual cell fate progression: cells on soft substrates, as a population, exhibit higher differentiation efficiency down the mesodermal lineage. Mesoderm differentiation has been shown to be affected by substrate mechanics in response to BMP4 and other soluble factors as well, including Activin A^30,67^. Other studies, using hPSCs cultured on conventional substrates such as glass or plastic, have reported that response to developmental signals can be affected by junctional integrity and myosin activity^68–70^. However, mechanosensing has, to our knowledge, not previously been connected to cellular competence for response to developmental signals and subsequent patterning. Here, we present a holistic cellular mechanism through which basement membrane mechanics influences hPSC epithelial organisation, which in turn directs cellular competence to regulate the spatial response to the fate-instructive BMP4 signal (Fig. 6e). In particular, reduced FAK-mediated mechanosensing leads to disruption of epithelial polarity and junctional integrity, which directly affect the accessibility of the otherwise basolaterally localised BMPR to the ligand. It is likely that the spatial response to other developmental signals whose receptors exhibit polarised localisation, including Wnt and Activin/Nodal^6,8^, could be affected by substrate stiffness changes through the same mechanism.

Our findings extend to the *in vivo* context of the mouse gastrulation showing that basement membrane mechanics also regulates the timing and pattern of fate specification in the mouse epiblast. It was previously shown that if collagen-rich basement membrane remodelling by MMPs is inhibited, mesodermal differentiation is reduced and gastrulation does not proceed^31^. We show that softening of the basement membrane by inhibiting collagen crosslinking is sufficient to induce premature mesoderm differentiation ectopically, throughout the epiblast and not just in the posterior midline as normally seen. This, in line with our *in vitro* data, suggests that mechanical signalling arising from the basement membrane contributes to regulation of both temporal and spatial response to developmental signals present in the pro-amniotic cavity. So far, it has been proposed that BMP4 diffuses through the gap between the extra- embryonic ectoderm and the epiblast to access basolateral BMPRs, which eventually causes a mesoderm differentiation wave in the epiblast^16^. However, this does not explain the spatiotemporal specificity of mesoderm induction. Based on our results, we hypothesise that epithelial reorganisation upon basement membrane softening initiated by increased MMP activity^31^ could lead to local and amplified response to BMP4 and mesoderm differentiation. It will also be interesting to investigate whether changes in epithelial polarity and permeability similarly contribute to enhancing response to signals in other contexts where the ECM is known to be remodelled, such as during cancer progression^71,72^.

Our data further show that apical ECM accumulation can modify cellular responses to soluble signals. hPSCs on soft substrates exhibit increased apical laminin accumulation, and strikingly, soft substrate-like BMP4 response can be mimicked in hPSCs on stiff substrates by supplying excess laminin to their apical surface. hPSCs have been recently reported to generate apical ECM flows using microvilli, influencing cell shape and gene expression^62^. Here, we have shown that culturing hPSCs on soft substrates alone leads to a higher density of microvilli than on stiff substrates, which could aid retention of ECM. Regardless of the underlying cause, the fact that apical ECM retention is sufficient to affect the cellular response to a soluble signal poses a new concept suggesting that apical ECM could condition spatiotemporal patterning. We also present evidence that presence of apical ECM on stiff substrates can reduce mechanosensing from the basement membrane, which suggests potentially critical feedback between mechanosensing and polarity. To our knowledge, this is the first evidence that the level of substrate mechanosensing can be overridden by supplying soluble ECM. Apical ECM is not very well studied, so whether apical ECM retention also contributes to changes in mechanosensing in other physiological and pathological states will be an interesting question for future studies.

Together, this study identifies a mechanism by which basement membrane mechanics shapes spatiotemporal responses to biochemical signals. Using the onset of gastrulation as an example, our findings underscore the importance of integrating mechanical and biochemical cues to enhance robustness of developmental patterning. These principles may also inform our understanding of disease progression, where ECM remodelling is a pervasive feature.

## MATERIALS AND METHODS

### Cell culture

hPSCs (Shef6 and H9 lines) used in this project were kind gifts from the Austin Smith lab. Cells were routinely cultured in TeSR-E8 medium (Stem Cell Technologies; #05990) on plastic plates coated with GelTrex (Thermo Scientific, #A1413301), in a humidified chamber at 37 °C and 5% CO2. The medium was exchanged every 24 h. Cells were passaged every 4-5 days using 0.5 EDTA (Thermo Scientific, #15-575-020) as a dissociation reagent for 3-4 minutes at 37 °C. Cells were passaged as clumps at a ratio of 1:12-1:16.

Before performing any experimental work involving hydrogels, cells were cultured on plastic plates coated with recombinant human vitronectin (10 μg/mL, Thermo Scientific; #A14700) in Dulbecco’s Phosphate Buffered Saline (PBS) (Sigma-Aldrich, #D8537) for 1 passage. The day before the experiment, cells were given mTeSR Plus medium (Stem Cell Technologies, #100-0276) and pre-treated with 10 μMof Y-27632/ROCK inhibitor (Stem Cell Technologies, #72302). During the experiments, cells were maintained in pure mTeSR Plus medium. Passage number never exceeded 50.

### StemBond hydrogel fabrication and functionalisation

StemBond hydrogels^25^ were fabricated by gel polymerisation between two glass coverslips. One coverslip (‘adherent coverslip’) was treated to adhere to the gel, while another (‘non-adherent coverslip’) was treated to be non-adherent for ease of detachment later. Adherent coverslips were first washed in 70% ethanol for 10 minutes, rinsed in distilled water and then washed in 0.2 M NaOH for 20 minutes. After another rinse in distilled water, coverslips were dried with Kimwipes and placed in Parafilm lined Petri dishes. Coverslips were then coated with a solution containing 5% 3- (Trimethoxysilyl)propyl methacrylate (Merck, #M6514) and 10% acetic acid diluted in ethanol for 2.5 h in a sealed chamber. Coverslips were subsequently cleaned by washing in ethanol and thoroughly wiped with ethanol-soaked Kimwipe to ensure removal of any remaining residues. Clean, adherent coverslips were placed face-up into clean Parafilm-coated Petri dishes. Non-adherent coverslips were washed in 70% ethanol for 20 minutes, rinsed in distilled water, and dried with Kimwipes. 2M AHA solution (IUPAC name: 2-Amino-6-(prop-2-enoylamino)hexanoic acid; AK Scientific, #6845AL) was prepared in methanol. This solution was used to prepare hydrogel solutions according to the recipes in Table 1. Polymerisation was initiated by adding 2.5 μl TEMED (Merck, #T22500) and 5 μl 10% APS (Merck, #A3678) to the solutions. The hydrogel mixture was pipetted immediately onto the adherent coverslip and subsequently covered with a non-adherent coverslip. Hydrogels were then left to polymerize in air for 45-50 minutes. The volume of hydrogel solution used depended on the coverslip size, but it was ensured that the thickness of the hydrogel remained greater than 100 μm. Finally, hydrogels were submerged in PBS containing 1% penicillin-streptomycin and stored at 4 °C. Non-adherent coverslips were removed from the hydrogels after overnight soaking in PBS by applying a small force with the tip of a scalpel inserted between the coverslips.

**Table 1.** Recipes for StemBond hydrogels with stiffnesses of 0.75, 3.5 and 160 kPa.

| **E (kPa)** | **Acrylamide 40% (μl)** | **Bis-acrylamide 2% (μl)** | **AHA 2 M (μl)** | **H<sub>2</sub>O (μl)** |
| --- | --- | --- | --- | --- |
| 0.75 | 35 | 27.2 | 20 | 410.3 |
| 3.5 | 48 | 36 | 20 | 388.5 |
| 160 | 200 | 150 | 20 | 122.5 |

To prepare for use, hydrogels were rinsed in MES/NaCl buffer (pH 6.1) and subsequently activated with 0.5M NHS (Acros Organics, #157272500) and 0.2M EDAC (Merck, #03450) in MES/NaCl buffer (pH 6.1) for 30 min. Hydrogels were then rinsed in pre-chilled PBS, followed by a rinse in 50 mM HEPES buffer (pH 8.5). They were subsequently coated with vitronectin (200 μg/mL; Thermo Scientific, #A14700) diluted in 50mM HEPES buffer (pH 8.5), overnight at 4 °C.

The next day, the coating medium was aspirated and hydrogels were blocked using 0.5M ethanolamine in HEPES buffer (pH 8.5) for 30 minutes. Finally, hydrogels were washed in PBS 3-4 times and stored in well plates containing DMEM F-12 with 1% penicillin-streptomycin (Thermo Scientific, #P4333). Before use, hydrogels were equilibrated in DMEM F-12 with 1% penicillin-streptomycin for 2 h at 37 °C after which the medium was changed to mTeSR Plus for another hour of equilibration at 37 °C. After dissociation with 0.5mM EDTA as described above, cells were seeded onto hydrogels submerged in mTeSR Plus.

### Pharmacological cell treatments

BMP4 (Bio-Techne, #314-BP-010) was administered to cells at a concentration of 50 ng/ml. Pharmacological treatments on the cells were performed as follows. FAK was inhibited using FAK Inhibitor 14 (10 μM; Abcam, #ab144503) for 1-1.5 h before BMP4 treatment.

PI3K inhibition was performed using LY294002 (50 μM; StemCell Technologies, #72152) for 2 h before BMP4 addition, at which point it was given again together with BMP4 (concentrations stated above) for another hour.

PTEN was inhibited using bisperoxovanadium (HOpic) (20 μM; Selleck Chemicals, #S8651) overnight before BMP4 treatment (concentrations as above).

Blebbistatin (Merck, #B0560) treatment was applied for 1 h at 100 μM.

Laminin treatment (Sigma-Aldrich, #L2020) was applied at a concentration of 50 μg/mL for 24 h before BMP4 addition.

In integrin activity inhibition experiments, on day 3 of culture, integrin β1 and integrin α6 were inhibited using CD29 (10 μg/mL; BioLegend, #102235) and CD49f (10 μg/mL; BioLegend, #313602) antibodies, respectively. The control samples were given Armenian Hamster IgG Isotype control antibody (10 μg/mL; BioLegend, #400969) and Purified Rat IgG2a, κ Isotype control antibody (10 μg/mL; BioLegend, #400501). Cells were pre-treated with the antibodies diluted in mTeSR Plus for 2 h before laminin was added to the medium at the concentration of 50 μg/mL. Antibodies and laminin were left on the cells overnight (at least 18 h). The next day, Medium was changed to mTeSR with BMP4 at a concentration of 50 ng/mL for 1 h before the cells were fixed.

### Immunostaining

#### General

Cells were fixed using 4% paraformaldehyde (PFA) for 20 min, and subsequently washed with PBS. Cells were then permeabilized with 0.25% Triton X-100 for 25-30 min (10 min in case of cell membrane proteins and no permeabilization for laminin). After washing with PBS, blocking was performed using 3% BSA in PBS for 1 h at room temperature. Primary antibody diluted in 1% BSA was added to the cells and left overnight at 4°C. The following primary antibodies were used: Brachyury (1:500; Cell Signalling Technology, #81694S), integrin β5 (1:200; Cell Signalling Technology, #3629S), pFAK (1:200; Thermo Scientific, #44-624G), active integrin β1 (1:100; Abcam, #ab30394), ZO-1 (1:250; Thermo Scientific, #33-9100), BMPR1A (1:50; Santa Cruz Biotechnology, #sc-518037), Myosin Regulatory Light Chain 2 (1:100; Cell Signalling Technology, #3672S), Claudin-6 (1:100; Santa Cruz Biotechnology, #sc-17669), laminin (1:200; Thermo Scientific, #PA1-16730). The next day, cells were washed in PBS before secondary antibody diluted in 1% BSA in PBS was added for 1 h at room temperature, with samples covered in foil. All secondary antibodies were Alexa Fluor purchased from Thermo Scientific and used at the concentration of 1:500. After PBS washes, samples were incubated in DAPI (Sigma-Aldrich, #D9542) and Phalloidin (Cell Signalling Technology, #8953) for 15 min and subsequently subjected to final PBS washes.

#### BMPR1A

BMPR1A was stained with the anti-BMPR1A antibody (Santa Cruz Biotechnology, #sc-518037) which was given to the live cells for 45 minutes at the concentration of 1:20 in mTeSR Plus before the cells were fixed. Cells were fixed with 4% PFA for 15 minutes and subsequently washed 3 times in PBS. No permeabilization was performed. The cells were subsequently incubated with the secondary antibody at a concentration of 1:500 in 1% BSA in PBS for 2 h at room temperature. Upon PBS washes, cells were treated with DAPI, Phalloidin and HCS CellMask for cytoplasm visualisation (Thermo Scientific, #H32721) for 30 minutes at room temperature. Cells were then thoroughly washed with PBS before imaging.

#### pSMAD1/5

Samples were fixed as described above. Subsequently, blocking and permeabilization were performed using 1% Triton X-100 and 3% BSA in PBS for 1 h. After a wash in PBS, samples were incubated in pre-warmed 1% SDS in PBS for 30 min at 37 °C. Samples were then washed 3 times in PBS. pSMAD1/5 antibody (1:800; Cell Signalling Technologies, #95165) was diluted in 1% Triton X-100 and 3% BSA in PBS. Samples were then incubated in a solution containing 1/3 of the described antibody solution and 2/3 of water, overnight at 4 °C. After washing in PBS, samples were incubated in the secondary antibody and DAPI diluted in 1% Triton X-100 and 3% BSA for 1.5 h at room temperature. Samples were subsequently washed in PBS.

#### Mouse embryos

Mouse embryos were fixed in 4% PFA for 30 min on a shaker at room temperature and subsequently washed 3 times with PBS. The embryos were permeabilised in 0.5% Triton-X in PBS for 30 min and subsequently blocked in 5% normal donkey serum, 0.3% BSA and 0.1% Triton-X in PBS for 2 hours. Primary antibodies were diluted in the blocking buffer and embryos were incubated overnight on a shaker at 4 °C. Antibodies were used at the following concentrations: T (1:100; Bio-techne, #AF2085), Collagen IV (1:100; Abcam, #ab19808). The next day, embryos were rinsed 3 times in 0.1% Triton-X in PBS before a longer wash in the same solution on a shaker for 20 min. Secondary antibodies were used at a concentration of 1:300, prepared in blocking buffer. Embryos were incubated in secondary antibodies for 1-2 h at room temperature. Subsequent washes in 0.1% Triton-X in PBS were performed before the embryos were stored in 0.1% Triton-X in PBS and 1:30 Vectashield Antifade Mounting Medium (Vector Laboratories, #H-1000-10).

### Imaging

#### Fluorescent microscopy

All fluorescent images were acquired using the confocal microscopes, Leica Sp8 and Stellaris 8. Where indicated, Structured Illumination Microscopy (SIM) was used to image BMPR and integrin β1.

#### Scanning electron microscopy

Prior to fixing, cells were rinsed three times with tissue culture-grade Dulbecco’s Phosphate Buffered Saline (PBS) (Sigma-Aldrich, #D8537). Samples were then fixed in 3% PFA, 2% glutaraldehyde in 0.05M sodiumcacodylate (pH 7.4) at room temperature for 1 h, and then overnight at 4 °C. Next, samples were thoroughly washed using distilled water 4 times. Subsequent dehydration was performed using increased concentrations of ethanol in the following sequence: 50% - 70% - 95% - 100% - 100%. Each step was done for 10 minutes on a gel shaker. Subsequently, samples were subjected to critical point drying and finally coated in gold with 8 nm thickness. Imaging was performed on the FEI Verios 460 scanning electron microscope at 2.00 kV and 25 pA.

### Transfections

#### Plasmid

To perform transfections, hPSCs were plated as single cells instead of clumps as is normally done when they are routinely passaged. For dissociation, cells were incubated in Accumax (Stem Cell Technologies, #07921) for 20 min at 37 °C. For 20mm hydrogels (12-well plate), 60,000 cells were seeded per well in mTeSR Plus supplemented with CloneR2 (StemCell Technologies, #100-0691).

18-24 h after seeding, the medium was exchanged to mTeSR Plus and cells were left to equilibrate in the newly added medium for 1 h. Transfection reagent was made by combining 6 μL of lipofectamine (Thermo Scientific; #STEM00003) with 3000 ng of AKT-PH-GFP plasmid DNA (Addgene, #18836) in 50 μL of mTeSR Plus. The mixture was added to one well containing 2 mL of mTeSR Plus. 24-30 h after transfection, medium was changed to pure mTeSR Plus. Cells were fixed 3 days after seeding.

#### siRNA

Cells were plated as single cells after a 20-minute incubation with Accumax (Stem Cell Technologies, #07921) at 37 °C. 300,000 cells were seeded per 35 mm diameter well in mTeSR Plus supplemented with CloneR2 (Stem Cell Technologies, #100-0691).

After 24 h, medium was changed to pure mTeSR Plus and siRNA was added at a final concentration of 25 nM. To achieve that, 9 μL of lipofectamine (Thermo Scientific, #13778100) was added to 150 μL of mTeSR Plus and 3 μL of siRNA (10 μM) was added to another 150 μL of mTeSR Plus. The two dilutions were mixed and 250 μL of the mix was added to 750 μL of pure mTeSR Plus in the well. siRNA was left on for 24 h before the medium was changed to pure mTeSR Plus and the cells were cultured for another 1-2 days.

CLDN6-targeting (Horizon Discovery, #L-015883-01-0005) and non-targeting (Horizon Discovery, #D-001810-10-05) siRNAs were purchased as SMARTPpools consisting of 4 different siRNAs.

### Assays

#### Western Blot

Cells grown on 6-well plates were scraped (2 wells per sample on hydrogels) and collected with 80 μL RIPA lysis buffer (Thermo Scientific, #89900) supplemented with phosphatase inhibitor (Roche, #04906845001) and protease inhibitor cocktail (Roche, #11836170001).

Samples were boiled for 5 minutes at 95 °C and subsequently sonicated for 10-15 cycles, 30 s on and 30 s off. Protein concentration was measured using the DC Protein Assay (Bio-Rad, #5000112) and the Pierce BCA Protein Assay (Thermo Scientific, #23227). 4x Laemmli buffer (Bio-Rad, #1610747) was diluted in the samples. Samples were boiled for 5minutes at 95 °C. 40 μg of protein per sample was loaded onto gels (Bio-Rad, #4561094) and run at 70 V for 15 minutes and then at 140 V. Trans-Blot TurboMini nitrocellulose membranes (Bio-Rad; #1704158) were used for transfer in the Bio-Rad transfer machine. Blocking was then performed using 1:1 PBS:Pierce StartingBlock Blocking Buffer (Thermo Scientific, #37538) for 1 h on a roller. Primary antibody diluted in the blocking buffer was left on the membrane overnight at 4 °C. The following primary antibodies were used: pFAK (1:1000; Thermo Scientific, #44-624G), pSMAD1/5 (1:1000; Cell Signalling Technology, #95165). Finally, the membrane was incubated in secondary antibody diluted in 5% BSA in TBST on a roller for 1 h at room temperature. Secondary antibodies were purchased from Licor and used at a concentration of 1:5000. Imaging of the membrane was performed on Licor Odyssey.

#### RNA extraction, cDNA synthesis and qPCR

RNA extraction was done using QiagenMini Plus Kit (Qiagen, #74134) following the manufacturer’s protocol. Subsequent cDNA synthesis was performed using the Superscript IV Reverse Transcriptase following the manufacturer’s instructions (Thermo Scientific, #18090010). RNA was subsequently degraded using Ribonuclease H (Thermo Scientific, #EN0201). qPCR was set up using the TaqMan Fast Universal PCRMasterMix (2x) (Applied Biosystems, #4352042) and TaqMan Gene Expression Assays. The following primers were used: GAPDH (Hs99999905_m1), T (Hs00610080_m1). Quantification was done on the StepOnePlus Real-Time PCR machine (Thermo Scientific) with GAPDH expression used as the endogenous control. Relative expression was calculated using the ΔΔCT method (Livak and Schmittgen, 2001).

### Dextran permeability experiment

To test permeability of the hPSC epithelium, fluorescein isothiocyanate–dextran of average molecular weight of 40 kDa (Sigma-Aldrich, #FD40S) was used. Cells were grown for 4 days on hydrogels polymerised on glass-bottom FluoroDish (WPI, #FD35-100). On day 4, cells were first incubated with CellMask (Thermo Scientific, #C10046) for 10-15 min to stain the cell membrane, and then washed twice with mTeSR Plus. Subsequently, cells were given mTeSR Plus supplemented with fluorescein isothiocyanate–dextran at the concentration of 50 μg/mL. Live-cell imaging was then immediately performed at 37 °C and 5% CO2 using the 40x silicon oil immersion objective (Olympus) on a spinning disk imaging system with a Perkin ElmerMLS1 base laser engine, CSU-X1 Spinning Disk head, Olympus IX81 stand, equipped with a Hamamatsu Flash 4.0v2 sCMOS camera and controlled by the Volocity software. Imaging was done across the height of the cells and a single sample was not imaged for longer than 1 h to match the standard BMP4 treatment length used across experiments.

### RNA Sequencing

Samples for RNA Sequencing were collected by pooling two 20 mm hydrogels. Cells were first dissociated by incubating in EDTA for 4 min at 37 °C. EDTA was then neutralised with cold mTeSR Plus. Cells were collected and spun down in 4 ml of mTeSR for 5 min at 300 x g. Supernatant was aspirated and the pellet was washed in cold PBS. PBS was subsequently aspirated and the pellet was snap-frozen at -80 °C. Each experiment was conducted on three separate occasions, and triplicates of each condition, representing biological replicates (N), were submitted to Azenta Life Sciences, where RNA Sequencing was performed using NovaSeq X + 2x150 bp platform.

Downstream analysis was also performed by the same company. Sequencing generated an average of ≈ 97 million paired-end reads per sample with high sequencing quality (mean Phred quality score of ≈ 38.6, with over 93% of bases having a Phred score of at least 30). Adapter trimming and quality filtering were performed using Trimmomatic v0.36. Reads were aligned to the human reference genome (GRCh38) using the STAR aligner v2.5.2b, which detects and incorporates splice junctions. Mapping rates exceeded 96% across all samples, with >92% of reads uniquely aligned. Gene-level quantification was conducted using feature Counts (Subread v1.5.2), counting only uniquely mapped reads overlapping exon features. Strand specificity was considered where applicable. Differential gene expression analysis was carried out using DESeq2 (DESeq2 1.36.0), applying the Wald test to calculate p-values and log2 fold changes. Genes with an adjusted p-value < 0.05 and an absolute log2 fold change > 1 were considered significantly differentially expressed.

The top two principal components were plotted in a PCA plot of the DESeq dataset following variance stabilising transformation.

Gene ontology enrichment analysis was performed using gseGEO (clusterProfiler_4.4.4), on all ontologies using ENSEMBL IDs (10,000 permutations, minimal gene set size 3, max 800, p-value cutoff of 0.05). Results were plotted using dotplot (enrichplot_1.16.2), splitting the sign of the normalised enrichment scores.

### Image quantifications

#### pSMAD1/5 signal intensity as a function of distance from colony edge

To quantify the pSMAD1/5 signal intensity as a function of distance from the colony edge, a custom script in Python was made for automated analysis. Images of pSMAD1/5 and DAPI staining acquired on the confocal microscope were used. Images which contained the entire extent of a colony were processed fully. If the colony size was bigger than the field of view, the colonies were imaged no further from the colony edge than the colony centre to minimise the chance that nuclei in the images were closer to colony edges outside of the field of view than edges inside the field of view. Nuclei were first segmented from the DAPI images using *CellPose*^73^. The pSMAD1/5 image with the corresponding segmented nuclei image were loaded into the script.

The purpose of the script was to assign each nucleus two measures – pSMAD1/5 signal intensity (y-axis) and distance to colony edge (x-axis). Firstly, the average pSMAD1/5 pixel value was calculated for each segmented nucleus. Secondly, to calculate the distance of each nucleus to the nearest colony edge, the centroid coordinates of each segmented nucleus were obtained using the “Measure region properties” function (scikit-image library). To obtain the distance of the nucleus centroid from the colony edge, firstly, the image of segmented nuclei was dilated and eroded using the binary closing function (SciPy library). This generated a mask of the colony with all pixels outside of the colony acquiring the “False” value. Using the distance transform function (SciPy library), an image was created where each pixel in the image was given a distance value to the nearest “False” value pixel, i.e., the nearest colony edge. Each nucleus centroid was hence given its corresponding distance value from the distance transformed image. Together, each nucleus then had a distance-to-edge value and a pSMAD1/5 average intensity value. Before generating the final plot, distances to colony edge were binned into 50 pixel-wide bins. The distance was subsequently converted from pixels to μm. Finally, mean and standard deviations of pSMAD1/5 intensities were plotted against their distance to colony edge.

#### Dextran penetration

To quantitatively compare the degree to which dextran penetrated lateral junctions as a measure of epithelial permeability, a custom image analysis script in Python for automated measurement was developed.

Cross-sectional (x-z) images of the cell monolayers were first generated from the original image stacks by re-slicing in Fiji. The CellMask images were then segmented in Tissue Analyzer to produce binary skeleton masks of cell edges (i.e., lateral junctions and apical and basal surfaces). Pixels belonging to cell interiors were identified from this mask by filling areas enclosed by the segmented cell edges (SciPy package). The mask was then parsed into edges and nodes using the Skan package. Edges representing lateral cell junctions were distinguished from apical and basal cell surfaces as those surrounded by cell interior pixels. These junctions were morphologically dilated to form the lateral- junction mask.

To ensure measurement of only dextran signal that had truly penetrated junctions, we excluded the regions of the lateral junction mask that were too close to the exterior of the cells. This was particularly important due to the noise caused by the strong dextran signal in the medium above the cells. We therefore used distance transforms (SciPy package) from the apical and basal edges to calculate, for each pixel in the cell interior, the pixel location as a fraction of the distance between the apical and basal surfaces. We could then keep just the central 50% of the lateral-junction mask (Extended Data Fig. 3d - green) by excluding pixels whose fraction was below 0.25 or above 0.75.

To measure the dextran signal intensity, we found that taking pixel intensity in these regions was unreliable due to variations in imaging conditions such as gel thickness and noise created by the strong dextran signal in the medium. Hence, we generated a thresholded signal by taking the top 4% (by dextran intensity) of pixels of the cell interior region (Extended Data Fig. 3d - red), reasoning that if dextran penetrated between the lateral junctions, we would find an enrichment of these pixels at the junctions. The final measure could then be defined as the fraction of the central lateral junction mask (Extended Data Fig. 3d - green) that was covered by thresholded signal pixels (Extended Data Fig. 3d - red).

#### Cell shape index

Outlines of cells were obtained through immunocytochemistry images of ZO-1. Images were processed using the Tissue Analyzer plugin in Fiji. This created a mask of the segmented cells which was analysed using a custom Python script. The sci-kit image package was used to measure the perimeter and area of the shapes of segmented cells. The cell shape index (CSI) was then calculated from the resulting measurements according to the following equation: 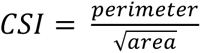.

#### BMPR1A/integrin β1 signal as a function of distance from cell surface

To quantify the amount of immunofluorescence signal as a function of distance from the apical surface of cell monolayers, cells were co-stained with CellMask HCS along with the antibody of interest. An automated image analysis pipeline was built in *Python*.

The pipeline uses the CellMask HCS channel to detect the apical surface in 3D confocal stacks. For this, the image was first lightly blurred to reduce noise. The width of the Gaussian used in the blur was chosen so that it was significantly smaller than the size of the smallest features appearing on the apical surface of the cells. A local sharpness measure was then calculated for each pixel, since this was expected to increase the signal-to-noise ratio of surface detection compared with fluorescence intensity alone. The algorithm first applied neighbourhood normalisation by dividing each pixel by the maximum value in a neighbour of defined size. The size here was chosen to be much larger than the typical size of the features that were being detected and ensured that the following steps were not influenced by regional variations in intensity.

The second step was to calculate the variance of the Laplacian of the 3D image to give the resulting sharpness measure. Two characteristic sizes were required here: the kernel sizes of the Laplacian and variance filters. For the Laplacian, a size larger than the typical distance over which an image object’s edge changed from a low to a high pixel value. For the variance kernel, a size roughly matching the size of a typical object in the image was chosen. A greyscale dilation, with the same kernel size as the earlier neighbourhood normalisation, was then applied. This has the effect of spreading the maximal pixel values in the X-Y plane, to “fill in” small regions of low sharpness resulting from the inhomogeneous distribution of features on the apical surface. Ultimately, this results in a smoother detected surface. Finally, to define the apical surface, this image was thresholded and the first true pixel across z-values for a given (x,y)-value was kept as "True", while the rest were changed to "False". The neighbourhood normalisation and sharpness measure of this algorithm allowed a single constant, 0.08, to be used as the thresholded value across all images.

A distance transform could then be applied to this apical surface mask (with values above the apical surface being made negative) to give the distance from the apical surface of every pixel. The final step was to extract a measure of signal as a function of this distance. To overcome the problem of a weak and sparse signal, which nevertheless had a clear spatial distribution, we used percentile thresholding, allowing us to analyse the spatial distribution of the brightest 0.1% percent of pixels in both conditions. Thus, the signal channel was thresholded to the 99.9th percentile pixel value and the fraction of thresholded pixels at each distance was plotted.

#### Apical/basolateral PIP3 ratio

To quantify the relative distribution of PIP3 in the membrane domains, we developed a *Python* script to calculate the ratio of apical and basolateral PH Akt-GFP, protein domain that binds directly to PIP3 in the membrane. In addition to imaging PH Akt-GFP, we also imaged the membrane stained with CellMask. These images were used to make two masks in Tissue Analyzer: the PH Akt-GFP images were used to segment the whole cell, while the CellMask image was used to segment the top the cells (apical cell surface). The latter mask was then used to split the whole-cell mask into the apical domain (where the two masks overlapped) and the basolateral domain (the rest of the whole-cell mask) (Extended Data Fig. 4b). Average signal intensity values were subsequently obtained from the PH Akt-GFP image for pixels corresponding to the apical domain mask and to the basolateral domain mask. The ratio of two values for individual cells was plotted as the apical/basolateral PIP3 distribution.

#### Intensity of pFAK focal points

To calculate the intensity of pFAK in focal points on the bottom of hPSC colonies, we developed a *Python* script for automated analysis. We denoised the images with a Gaussian blur and calculated the multi-Otsu thresholds with 3 classes (Sci-kit learn package). The 3 classes were used to represent the 3 groups of pixel intensities found in the image: background, cell background and signal. We took the brightest class of pixels (signal) to be the true pFAK signal. Using this threshold, were produced masks of pFAK focal points. Masks were subsequently used to extract the intensities of pixels of pFAK signal in focal points.

### Mouse embryo dissections and culture

Female (4-8 weeks old) and male (> 8 weeks old) CD1 mice were time-mated under standard housing conditions (12-h light–dark cycle, ambient temperature 19–22 °C, humidity 45–65%). Mouse embryos were dissected at either E6.5 or E5.5 from the maternal decidua in DMEM F-12 medium (Sigma-Aldrich, #D6421) supplemented with 5% FBS, with intact Reichert’s membrane and ectoplacental cone. The sex of the embryos was not determined. The dissected E6.5 embryos were fixed immediately in 4% PFA while E5.5 embryos were cultured in DMEM/F-12 supplemented with Glutamax (ThermoScientific, #35050061) and 50% rat serum (Charles River Laboratories) at 37 °C and 5% CO2. Littermate embryos were split between control and treatment conditions. Inhibition of Collagen IV crosslinking in culture was performed using β Aminopropionitrile (BAPN) (Sigma-Aldrich, #A3134) at a concentration of 500 μM– 1mM. After 16 h of culture, control and treated embryos were washed briefly in PBS and fixed in 4% PFA for 30 minutes at room temperature. Embryo numbers were pooled from three individually conducted experiments, representing biological replicates (N).

## ACKNOWLEDGEMENTS

We thank Daniel St Johnston for comments on the manuscript, Karin Mueller and Filomena Gallo (Cambridge Advanced Imaging Centre) for SEM sample preparation, Darran Clements (Cambridge Stem Cell Institute Imaging Core Facility) for assistance in microscopy, Jeanne Lefévère-Laoide for assistance with SIM, and Fiona Morgan and Oliver Bower for technical support. This work was supported by the Wellcome Trust (218481/Z/19/Z to A.R. and 221856/Z/20/Z to K.K.N.), a European Research Council Consolidator Grant (772798 to K.J.C.), and a core support grant from the Wellcome Trust (203151/Z/16/Z, 203151/A/16/Z) and the UKRI Medical Research Council (MC_PC_17230) to the Cambridge Stem Cell Institute.

## AUTHOR CONTRIBUTIONS

K.J.C., A.R and E.K.P designed the study. K.J.C. and E.K.P supervised the research. A.R., K.J.C and E.K.P. wrote the manuscript. A.R. performed all hPSC experiments. T.P.J.W. and A.R. wrote codes for image analysis. C.S.S. and A.R. performed the mouse embryo experiments. K.K.N. supervised the mouse embryo experiment conception. L.M.D.W. carried out the bulk RNA sequencing analysis. All authors discussed the manuscript.

## EXTENDED DATA

**Extended Data Figure 1.**
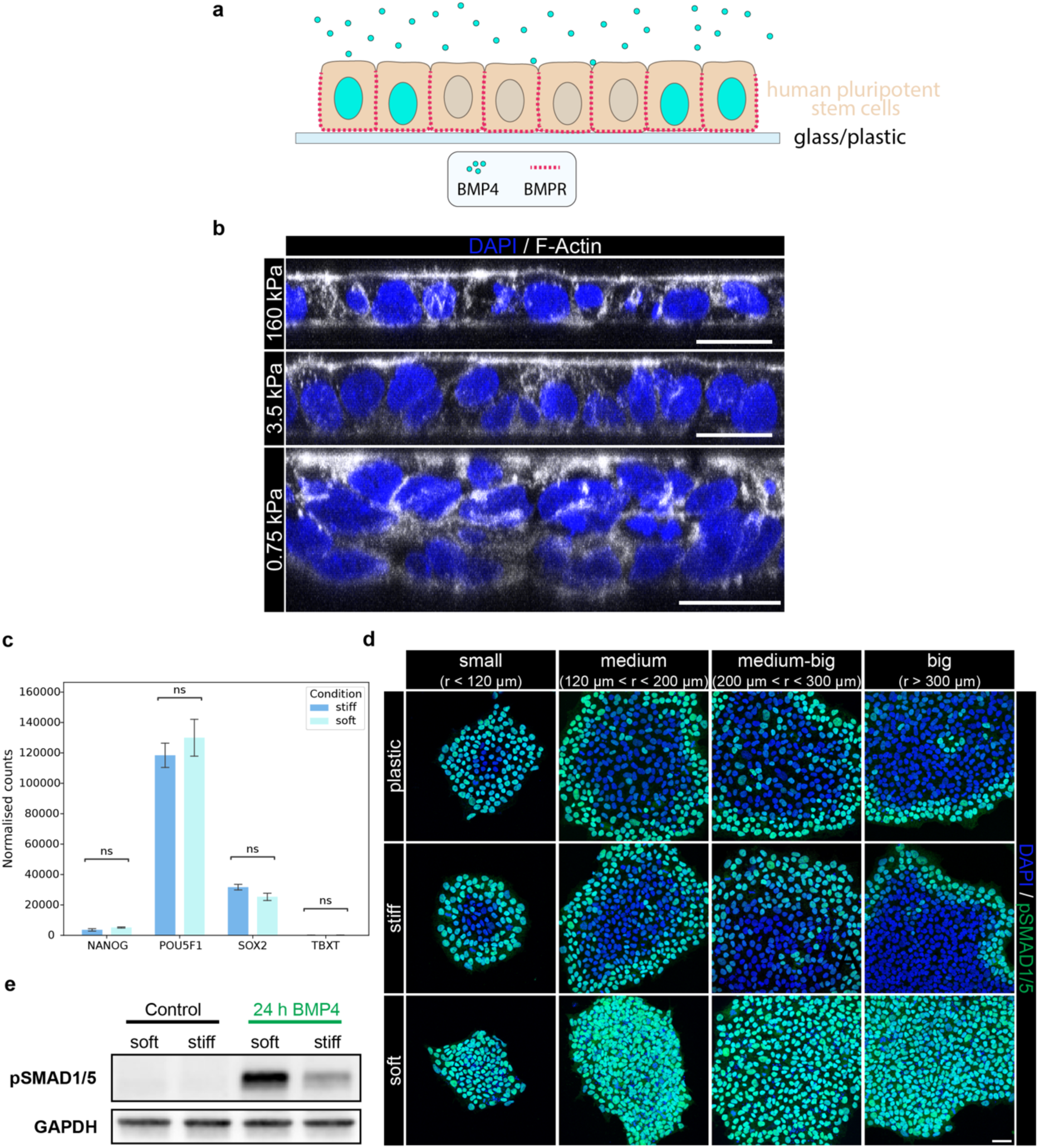
Substrate stiffness regulates the response of hPSCs to BMP4 but alone does not induce differentiation, related to Figure 1. a. Schematic representing the known response of hPSCs on glass/plastic to 1 h of uniform apical exposure to the BMP4 signal. Such response has been described to be conditioned by the basolateral localisation of the BMP receptors (BMPRs) and high junctional integrity. b. Representative immunofluorescence merged images (maximum projection) of hPSCs on hydrogels of various stiffnesses, stained for F-Actin and DAPI. Images were reconstructed in x-z plane; 10 images were projected in y-axis. Scale bar = 20 μm. c. Normalised counts obtained from bulk RNA-sequencing data of hPSCs on stiff and soft substrates for pluripotency markers (*NANOG, POUF51, SOX2*) and *TBXT* (Brachyury). Differential expression analysis was performed using DESeq2 with the Wald test. Data was generated from N = 3 independent experiments representing biological replicates. d. Representative immunofluorescence merged images (maximum projection) of hPSCs on plastic, stiff and soft substrates, treated with BMP4 for 1 h and stained for pSMAD1/5 and DAPI. Scale bar = 50 μm. Individual data came from N = 3 independent experiments representing biological replicates. e. Western Blot of pSMAD1/5 and GAPDH from hPSCs on stiff and soft substrates without BMP4 treatment (control) and after 24 h of BMP4 treatment.

**Extended Data Figure 2.**
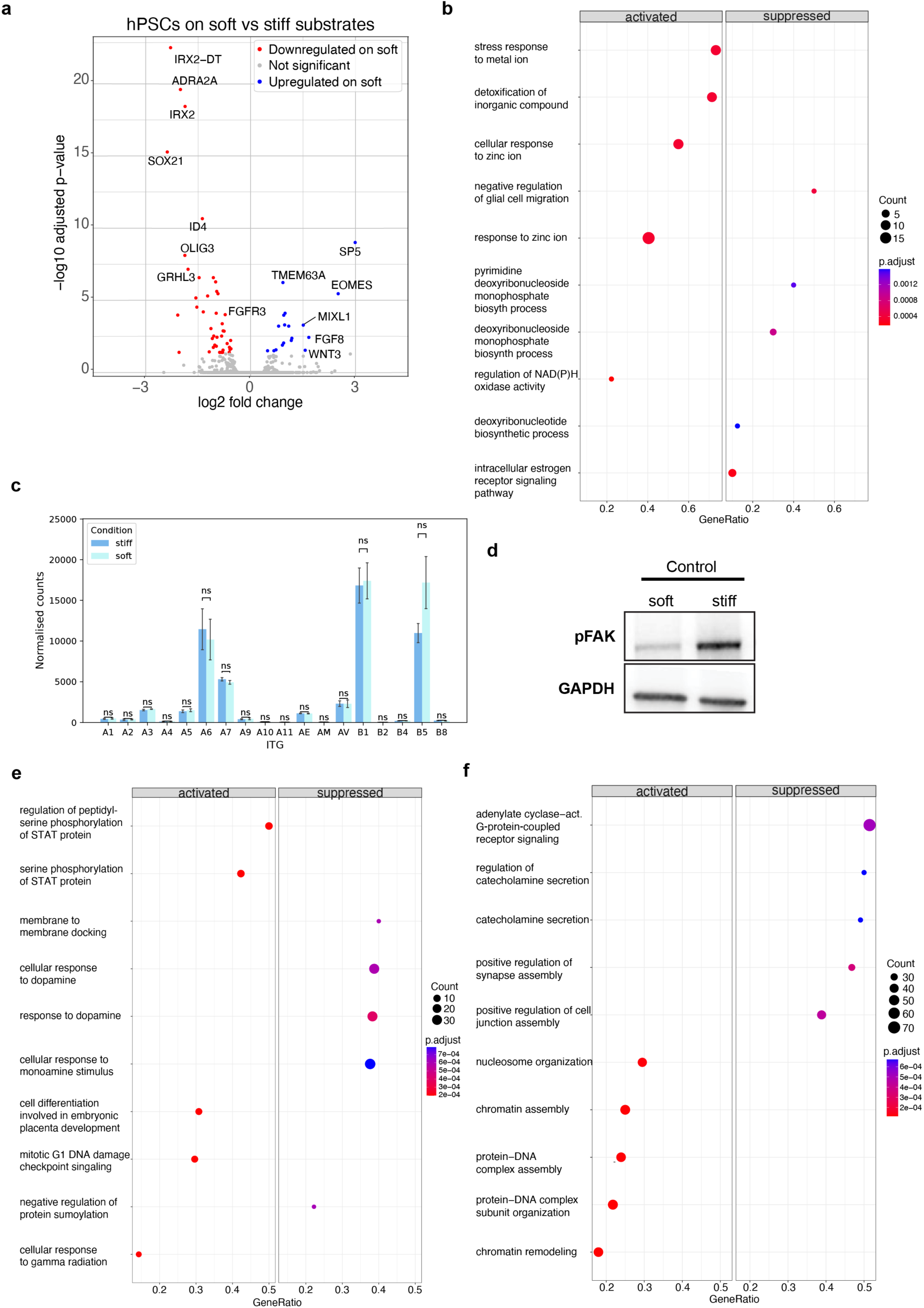
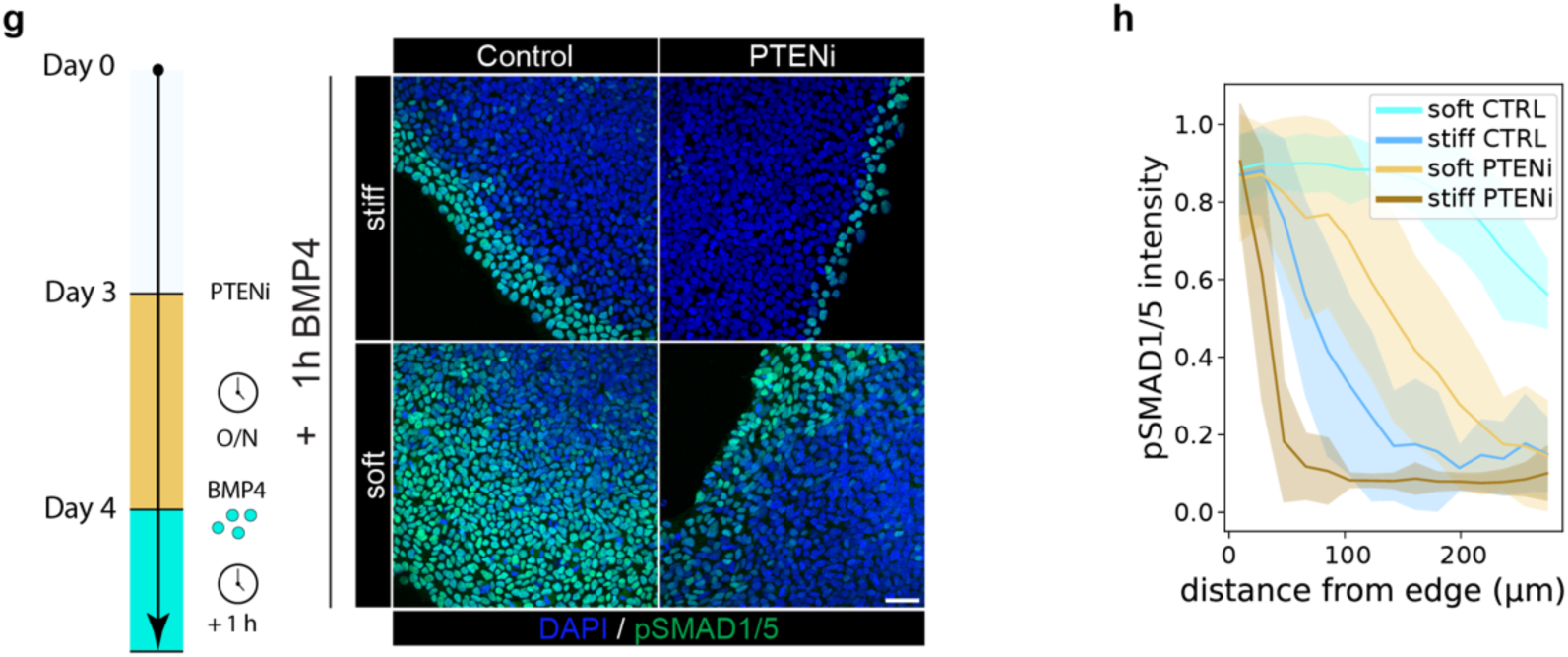
Transcriptional profiles for hPSCs on different substrate stiffnesses and the effects of perturbation of the FAK-PI3K cascade, related to Figure 2. a. Volcano plot of DEGs showing the log2 fold change (x axis) of hPSCs on soft substrates relative to stiff substrates as a function of the -log10(adjusted p value) (y axis). In blue are genes that are significantly upregulated on soft substrates at a p-adjusted value of 0.05. In red are genes that are significantly downregulated on soft substrates at a p-adjusted value of 0.05. The total number of significantly differentially expressed genes equals 62. Data was generated from N = 3 independent experiments representing biological replicates. b. Dot plot of enriched GO terms for the soft control vs stiff control comparison. The top 5 GO biological processes with the largest gene ratios are shown. The size of the dots represents the number of genes in the significant DEG list associated with the GO term and the colour represents the p-adjusted values. The plot is split by sign of the normalised enrichment score (NES, NES>0 as activated, NES<0 as suppressed). Data was generated from N = 3 independent experiments representing biological replicates. c. Normalised counts obtained from RNA-sequencing data of hPSCs on stiff and soft substrates for each integrin (*ITG*) subunit expressed. Differential expression analysis was performed using DESeq2 with the Wald test. Data was generated from N = 3 independent experiments representing biological replicates. d. Western Blot of pFAK and GAPDH in hPSCs on stiff and soft substrates without BMP4 treatment (control). e. Dot plot of enriched GO terms for the stiff FAK inhibitor vs stiff control comparison. The top 5 GO biological processes with the largest gene ratios are shown. The size of the dots represents the number of genes in the significant DEG list associated with the GO term and the colour represents the p-adjusted values. The plot is split by sign of the normalised enrichment score (NES, NES>0 as activated, NES<0 as suppressed). Data was generated from N = 3 independent experiments representing biological replicates. from N = 3 independent experiments representing biological replicates. f. Dot plot of enriched GO terms for the stiff PI3K inhibitor vs stiff control comparison. The top 5 GO biological processes with the largest gene ratios are shown. The size of the dots represents the number of genes in the significant DEG list associated with the GO term and the colour represents the p-adjusted values. The plot is split by sign of the normalised enrichment score (NES, NES>0 as activated, NES<0 as suppressed). Data was generated from N = 3 independent experiments representing biological replicates. g. Left: Schematic representing the experimental protocol involving PTEN inhibition (PTENi) before BMP4 treatment. Right: Representative immunofluorescence merged images (maximum projection) of hPSCs on stiff and soft substrates, treated with BMP4 for 1 h, with or without prior PTEN inhibition, stained for pSMAD1/5 and DAPI. Scale bar = 50 μm. h. Quantification of pSMAD1/5 signal from immunofluorescence images of hPSCs in stiff and soft control, and stiff and soft PTENi condition, after 1 h of BMP4 treatment. The signal intensity in each nucleus is plotted as a function of the distance of that nucleus from the colony edge, normalized by the 98^th^ percentile value of each image. The continuous line represents the mean signal intensity and edges of the shaded areas represent +/- standard deviation from N = 3 biological replicates.

**Extended Data Figure 3.**
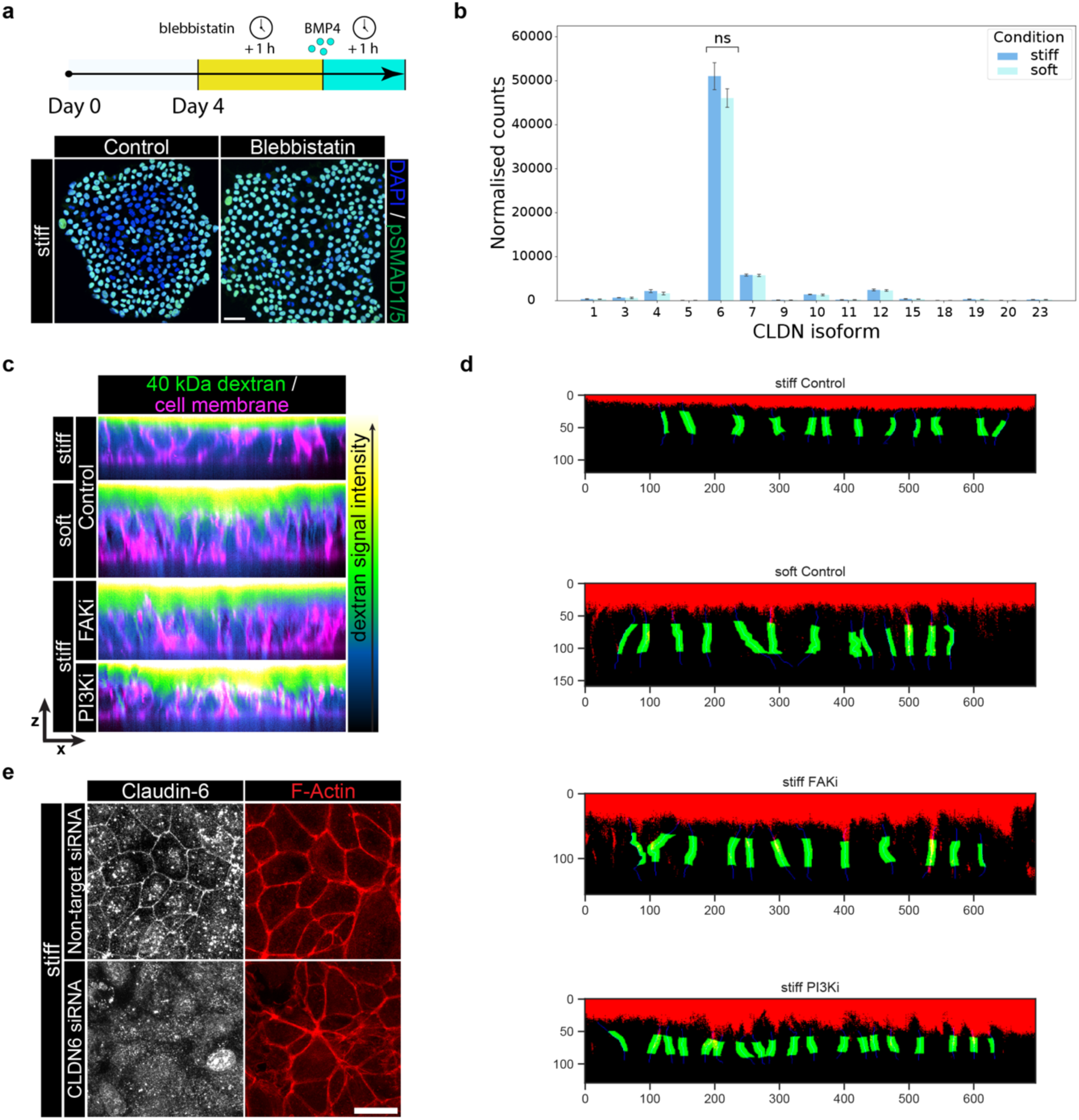
Supplementary data on junctional integrity reduction and increased permeability on soft substrates, related to Figure 3. a. Top: Schematic representing the experimental protocol involving myosin activity inhibition using blebbistatin before the BMP4 treatment. Bottom: Representative immunofluorescence merged images (maximum projection) of hPSCs on stiff substrates with (blebbistatin) or without (control) myosin activity inhibition and after 1 h of BMP4 treatment, stained for pSMAD1/5 and DAPI. Scale bar = 50 μm. b. Normalised counts obtained from RNA-sequencing data of hPSCs on stiff and soft substrates for each Claudin (*CLDN*) gene expressed. Differential expression analysis was performed using DESeq2 with the Wald test. Data was generated from N = 3 independent experiments representing biological replicates. c. Representative merged images of fluorescent dextran signal and cell membrane dye (CellMask) along the apicobasal axis (x-z plane) on stiff and soft substrates, in the control condition or after FAK inhibitor (FAKi) or PI3K inhibitor (PI3Ki) treatment. The lookup table indicates dextran signal intensity from low (blue) to high (white). d. Images produced after processing dextran images and cell membrane images in *Python*. These images were used for quantification of dextran signal at the lateral cell junctions. Dextran images were thresholded by keeping the top 4% of brightest pixels (red). All other pixels in the image were set to 0. Blue lines represent masks of lateral junctions generated from the cell membrane staining. Green area represents the morphologically dilated, central 50% of the lateral junction mask. e. Representative immunofluorescence images (maximum projection) of colony top (top 5 slices) of hPSCs on stiff substrates, treated with Claudin-6 targeting siRNAs or non-targeting (control) siRNAs, stained for Claudin-6 or F-Actin. Scale bar = 20 μm. Individual data came from N = 3 independent experiments representing biological replicates.

**Extended Data Figure 4.**
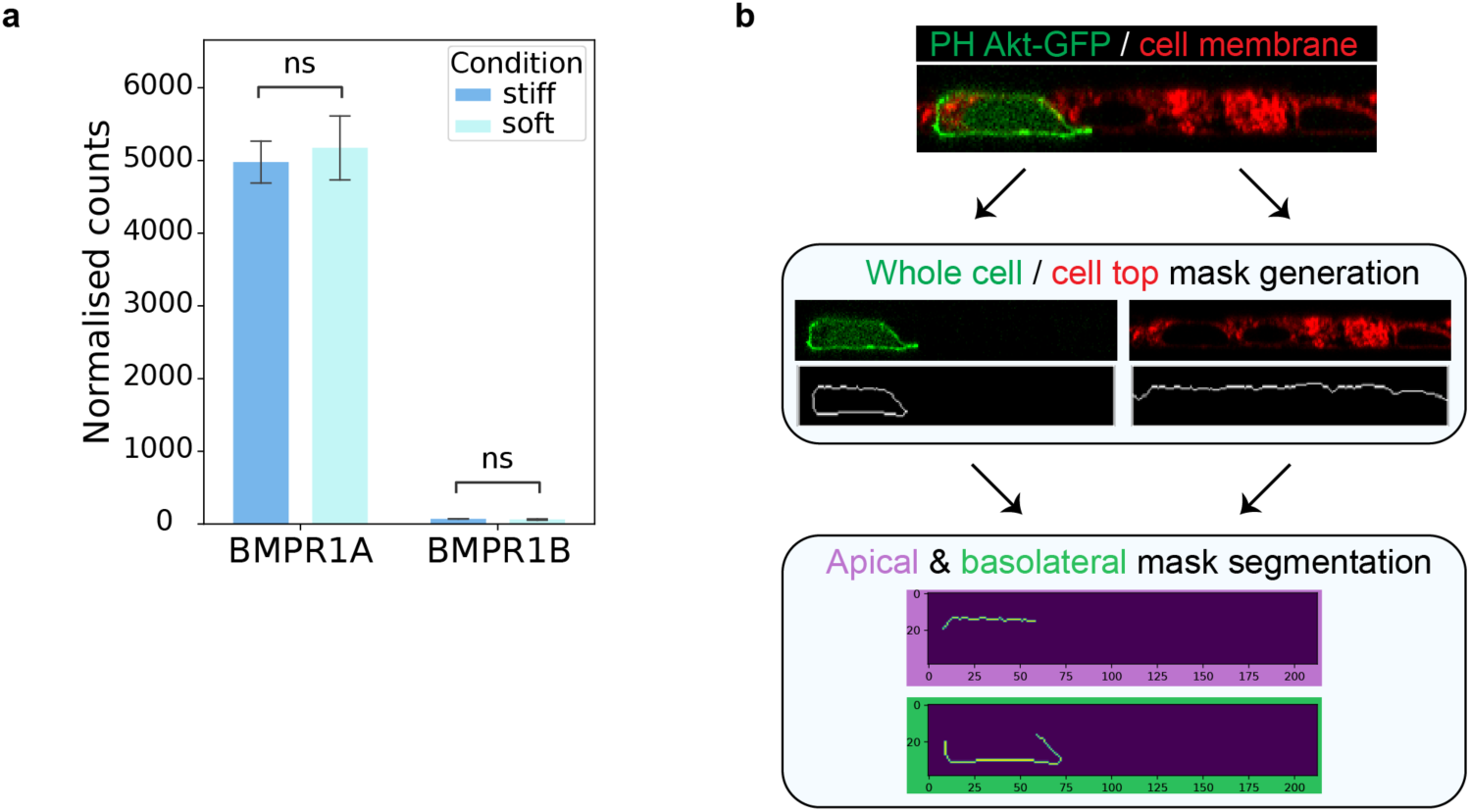
Additional information for the quantifications of apico-basal distributions in hPSCs, related to Figure 4. a. Normalised counts obtained from RNA-sequencing data of hPSCs on stiff and soft substrates for *BMPR1A* and *BMPR1B*. Differential expression analysis was performed using DESeq2 with the Wald test. Data was generated from N = 3 independent experiments representing biological replicates. b. Pipeline used for quantification of the apical/basolateral ratio of PH Akt-GFP (labelling PIP3) signal intensity. See *Materials and Methods*.

**Extended Data Figure 5.**
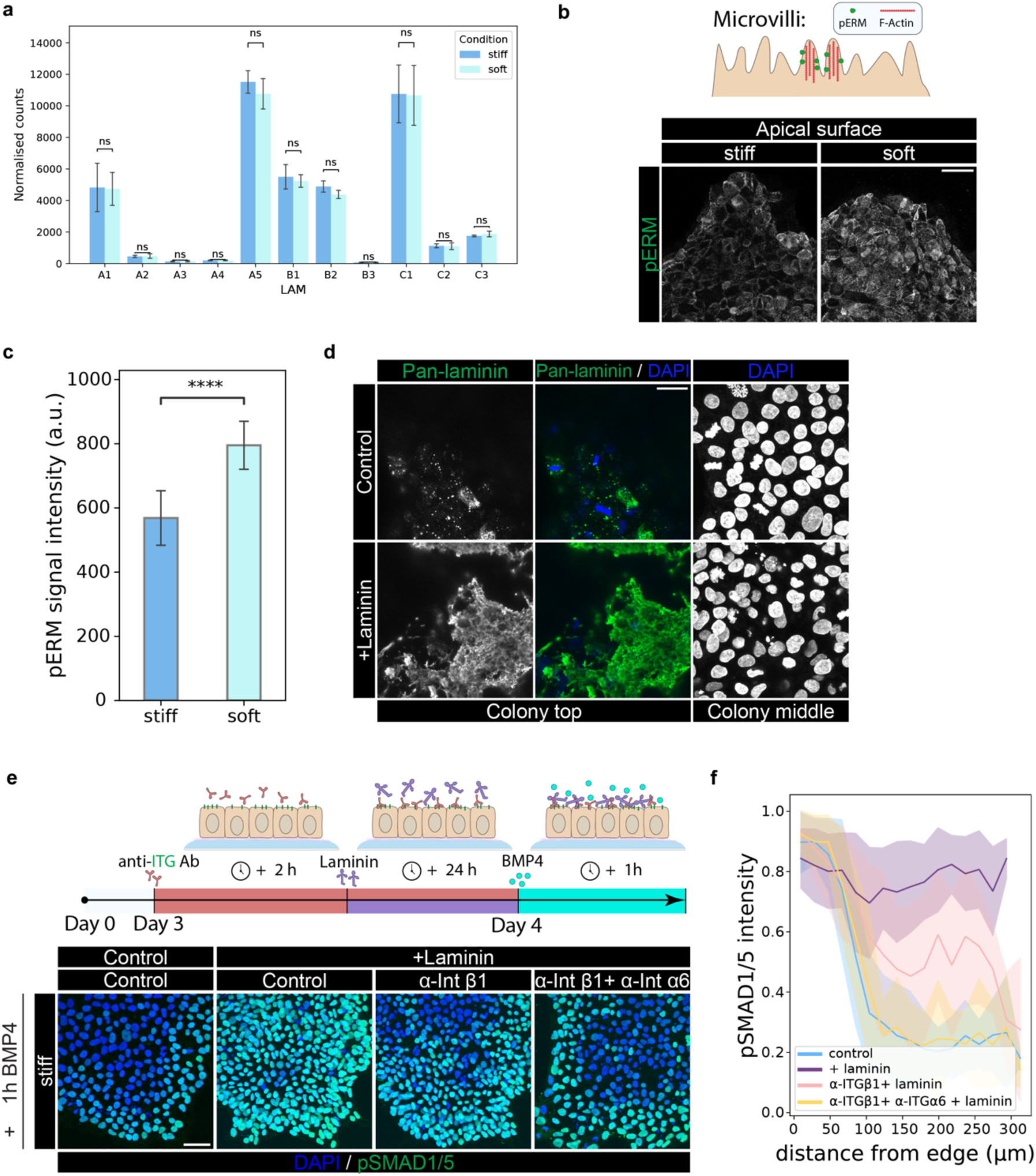
Apical laminin accumulation and subsequent integrin activation disrupts patterned response to BMP4, related to Figure 5. a. Normalised counts obtained from RNA-sequencing data of hPSCs on stiff and soft substrates for each laminin (*LAM*) subunit expressed. Differential expression analysis was performed using DESeq2 with the Wald test. Data was generated from N = 3 independent experiments representing biological replicates. b. Top: Schematic depicting microvilli architecture. Bottom: Representative immunofluorescence images (maximum projection) of apical surfaces (top 10 slices) of hPSCs on stiff and soft substrates, stained for pERM. Scale bar = 50 μm. c. Quantification of pERM signal intensity across maximum projection images. Individual data came from N = 3 independent experiments representing biological replicates. P-value determined by Welch’s t-test. **** indicates p-value < 0.0001. d. Representative immunofluorescence images of apical colony surface of hPSCs on stiff and soft substrates, stained for laminin and DAPI, and colony middle stained for DAPI. Scale bar = 25 μm. e. Top: Schematic representing the experimental protocol involving inhibition of integrin activity with antibodies, prior to soluble laminin addition into the cell medium and BMP4 treatment. Bottom: Representative immunofluorescence merged images (maximum projection) of hPSCs on stiff substrates stained for pSMAD1/5 and DAPI after 1 h of BMP4 treatment, treated or untreated with soluble laminin and treated or untreated with the anti-integrin β1 and anti-integrin α6 antibodies. Scale bar = 50 μm. f. Quantification of normalised pSMAD1/5 signal intensity from immunofluorescence images of hPSCs on stiff and soft substrates after 1 h of BMP4 treatment in the control condition (blue), upon addition of soluble laminin (purple) and after inhibition of integrin β1 (pink) and both integrin β1 and integrin α6 inhibition (yellow) before soluble laminin addition. The signal intensity in each nucleus is plotted as a function of the distance of that nucleus from the colony edge. The signal intensity is normalized to the 98^th^ percentile. The continuous line represents the mean signal intensity and edges of the shaded areas represent +/- standard deviation from N = 3 biological replicates.

## REFERENCES

1. Turing, A. M. The chemical basis of morphogenesis. Philos. Trans. R. Soc. Lond. B. Biol. Sci. 237, 37–72 (1952).

2. Wolpert, L. Positional information and the spatial pattern of cellular differentiation. J. Theor. Biol. 25, 1–47 (1969).

3. Rogers, K. W. & Schier, A. F. Morphogen gradients: from generation to interpretation. Annu. Rev. Cell Dev. Biol. 27, 377–407 (2011).

4. Kondo, S., Iwashita, M. & Yamaguchi, M. How animals get their skin patterns: fish pigment pattern as a live Turing wave. Int. J. Dev. Biol. 53, 851–856 (2009).

5. Raspopovic, J., Marcon, L., Russo, L. & Sharpe, J. Modeling digits. Digit patterning is controlled by a Bmp-Sox9-Wnt Turing network modulated by morphogen gradients. Science 345, 566–570 (2014).

6. Etoc, F. et al. A Balance between Secreted Inhibitors and Edge Sensing Controls Gastruloid Self-Organization. Dev. Cell **39**, 302–315 (2016).

7. Simunovic, M. et al. A 3D model of a human epiblast reveals BMP4-driven symmetry breaking. Nat. Cell Biol. 21, 900–910 (2019).

8. Martyn, I., Brivanlou, A. H. & Siggia, E. D. A wave of WNT signaling balanced by secreted inhibitors controls primitive streak formation in micropattern colonies of human embryonic stem cells. Development 146, dev172791 (2019).

9. Huang, Y.-L. & Niehrs, C. Polarized Wnt Signaling Regulates Ectodermal Cell Fate in *Xenopus*. Dev. Cell **29**, 250–257 (2014).

10. Gui, J., Huang, Y. & Shimmi, O. Scribbled Optimizes BMP Signaling through Its Receptor Internalization to the Rab5 Endosome and Promote Robust Epithelial Morphogenesis. PLOS Genet. 12, e1006424 (2016).

11. Winnier, G., Blessing, M., Labosky, P. A. & Hogan, B. L. Bone morphogenetic protein-4 is required for mesoderm formation and patterning in the mouse. Genes Dev. 9, 2105–2116 (1995).

12. Ben-Haim, N. et al. The nodal precursor acting via activin receptors induces mesoderm by maintaining a source of its convertases and BMP4. Dev. Cell 11, 313– 323 (2006).

13. Arnold, S. J. & Robertson, E. J. Making a commitment: cell lineage allocation and axis patterning in the early mouse embryo. Nat. Rev. Mol. Cell Biol. 10, 91–103 (2009).

14. Perea-Gomez, A. et al. Nodal antagonists in the anterior visceral endoderm prevent the formation of multiple primitive streaks. Dev. Cell 3, 745–756 (2002).

15. Thowfeequ, S. et al. An integrated approach identifies the molecular underpinnings of murine anterior visceral endoderm migration. Dev. Cell **59**, 2347–2363.e9 (2024).

16. Zhang, Z., Zwick, S., Loew, E., Grimley, J. S. & Ramanathan, S. Mouse embryo geometry drives formation of robust signaling gradients through receptor localization. Nat. Commun. 10, 4516 (2019).

17. Nakamura, T. et al. A developmental coordinate of pluripotency among mice, monkeys and humans. Nature 537, 57–62 (2016).

18. Rossant, J. & Tam, P. P. L. New Insights into Early Human Development: Lessons for Stem Cell Derivation and Differentiation. Cell Stem Cell 20, 18–28 (2017).

19. Warmflash, A., Sorre, B., Etoc, F., Siggia, E. D. & Brivanlou, A. H. A method to recapitulate early embryonic spatial patterning in human embryonic stem cells. Nat. Methods 11, 847–854 (2014).

20. Minn, K. T. et al. High-resolution transcriptional and morphogenetic profiling of cells from micropatterned human ESC gastruloid cultures. eLife **9**, e59445 (2020).

21. Engler, A. J., Sen, S., Sweeney, H. L. & Discher, D. E. Matrix elasticity directs stem cell lineage specification. Cell 126, 677–689 (2006).

22. Huebsch, N. et al. Harnessing traction-mediated manipulation of the cell/matrix interface to control stem-cell fate. Nat. Mater. **9**, 518–526 (2010).

23. Segel, M. et al. Niche stiffness underlies the ageing of central nervous system progenitor cells. Nature 573, 130–134 (2019).

24. Chowdhury, F. et al. Material properties of the cell dictate stress-induced spreading and differentiation in embryonic stem cells. Nat. Mater. **9**, 82–88 (2010).

25. Labouesse, C. et al. StemBond hydrogels control the mechanical microenvironment for pluripotent stem cells. Nat. Commun. 12, 6132 (2021).

26. Rowlands, A. S., George, P. A. & Cooper-White, J. J. Directing osteogenic and myogenic differentiation of MSCs: interplay of stiffness and adhesive ligand presentation. Am. J. Physiol. Cell Physiol. 295, C1037–1044 (2008).

27. Khetan, S. et al. Degradation-mediated cellular traction directs stem cell fate in covalently crosslinked three-dimensional hydrogels. Nat. Mater. 12, 458–465 (2013).

28. Guvendiren, M. & Burdick, J. A. The control of stem cell morphology and differentiation by hydrogel surface wrinkles. Biomaterials 31, 6511–6518 (2010).

29. Kilian, K. A., Bugarija, B., Lahn, B. T. & Mrksich, M. Geometric cues for directing the differentiation of mesenchymal stem cells. Proc. Natl. Acad. Sci. U. S. A. 107, 4872– 4877 (2010).

30. Przybyla, L., Lakins, J. N. & Weaver, V. M. Tissue Mechanics Orchestrate Wnt-Dependent Human Embryonic Stem Cell Differentiation. Cell Stem Cell **19**, 462–475 (2016).

31. Kyprianou, C. et al. Basement membrane remodelling regulates mouse embryogenesis. Nature 582, 253–258 (2020).

32. Chen, D.-Y. et al. Basement membrane perforations guide anterior–posterior axis formation. Nat. Commun. 16, 6763 (2025).

33. Wilkinson, D. G., Bhatt, S. & Herrmann, B. G. Expression pattern of the mouse T gene and its role in mesoderm formation. Nature 343, 657–659 (1990).

34. Miyazono, K., Kamiya, Y. & Morikawa, M. Bone morphogenetic protein receptors and signal transduction. J. Biochem. (Tokyo) 147, 35–51 (2010).

35. Chhabra, S., Liu, L., Goh, R., Kong, X. & Warmflash, A. Dissecting the dynamics of signaling events in the BMP, WNT, and NODAL cascade during self-organized fate patterning in human gastruloids. PLOS Biol. 17, e3000498 (2019).

36. Kechagia, J. Z., Ivaska, J. & Roca-Cusachs, P. Integrins as biomechanical sensors of the microenvironment. Nat. Rev. Mol. Cell Biol. 20, 457–473 (2019).

37. Iwamoto, D. V. & Calderwood, D. A. Regulation of integrin-mediated adhesions. Curr. Opin. Cell Biol. 36, 41–47 (2015).

38. Kornberg, L., Earp, H. S., Parsons, J. T., Schaller, M. & Juliano, R. L. Cell adhesion or integrin clustering increases phosphorylation of a focal adhesion-associated tyrosine kinase. J. Biol. Chem. 267, 23439–23442 (1992).

39. Kanchanawong, P. & Calderwood, D. A. Organization, dynamics and mechanoregulation of integrin-mediated cell–ECM adhesions. Nat. Rev. Mol. Cell Biol. 24, 142–161 (2023).

40. Schlaepfer, D. D., Hauck, C. R. & Sieg, D. J. Signaling through focal adhesion kinase. Prog. Biophys. Mol. Biol. 71, 435–478 (1999).

41. Golubovskaya, V. M. et al. A small molecule inhibitor 1,2,4,5-Benzenetetraamine tetrahydrochloride, targeting the Y397 site of Focal Adhesion Kinase decreases tumor growth. J. Med. Chem. 51, 7405–7416 (2008).

42. Chen, H. C., Appeddu, P. A., Isoda, H. & Guan, J. L. Phosphorylation of tyrosine 397 in focal adhesion kinase is required for binding phosphatidylinositol 3-kinase. J. Biol. Chem. 271, 26329–26334 (1996).

43. Vlahos, C. J., Matter, W. F., Hui, K. Y. & Brown, R. F. A specific inhibitor of phosphatidylinositol 3-kinase, 2-(4-morpholinyl)-8-phenyl-4H-1-benzopyran-4-one (LY294002). J. Biol. Chem. 269, 5241–5248 (1994).

44. Maehama, T. & Dixon, J. E. The tumor suppressor, PTEN/MMAC1, dephosphorylates the lipid second messenger, phosphatidylinositol 3,4,5-trisphosphate. J. Biol. Chem. 273, 13375–13378 (1998).

45. Stambolic, V. et al. Negative regulation of PKB/Akt-dependent cell survival by the tumor suppressor PTEN. Cell **95**, 29–39 (1998).

46. Rodriguez-Boulan, E. & Macara, I. G. Organization and execution of the epithelial polarity programme. Nat. Rev. Mol. Cell Biol. 15, 225–242 (2014).

47. Saitoh, M. et al. Basolateral BMP Signaling in Polarized Epithelial Cells. PLoS ONE 8, e62659 (2013).

48. Suzuki, T. Regulation of intestinal epithelial permeability by tight junctions. Cell. Mol. Life Sci. CMLS 70, 631–659 (2013).

49. Quiros, M. & Nusrat, A. RhoGTPases, actomyosin signaling and regulation of the Epithelial Apical Junctional Complex. Semin. Cell Dev. Biol. 36, 194–203 (2014).

50. Ivanov, A. I. et al. A unique role for nonmuscle myosin heavy chain IIA in regulation of epithelial apical junctions. PloS One 2, e658 (2007).

51. Naydenov, N. G. et al. Nonmuscle Myosin IIA Regulates Intestinal Epithelial Barrier in vivo and Plays a Protective Role During Experimental Colitis. Sci. Rep. 6, 24161 (2016).

52. Belardi, B. et al. A Weak Link with Actin Organizes Tight Junctions to Control Epithelial Permeability. Dev. Cell 54, 792–804.e7 (2020).

53. Lynn, K. S., Peterson, R. J. & Koval, M. Ruffles and spikes: Control of tight junction morphology and permeability by claudins. Biochim. Biophys. Acta BBA - Biomembr. 1862, 183339 (2020).

54. Ben-David, U., Nudel, N. & Benvenisty, N. Immunologic and chemical targeting of the tight-junction protein Claudin-6 eliminates tumorigenic human pluripotent stem cells. Nat. Commun. 4, 1992 (2013).

55. Buckley, C. E. & St Johnston, D. Apical-basal polarity and the control of epithelial form and function. Nat. Rev. Mol. Cell Biol. 23, 559–577 (2022).

56. Gassama-Diagne, A. et al. Phosphatidylinositol-3,4,5-trisphosphate regulates the formation of the basolateral plasma membrane in epithelial cells. Nat. Cell Biol. 8, 963–970 (2006).

57. Martin-Belmonte, F. et al. PTEN-mediated apical segregation of phosphoinositides controls epithelial morphogenesis through Cdc42. Cell 128, 383–397 (2007).

58. Lee, J. L. & Streuli, C. H. Integrins and epithelial cell polarity. J. Cell Sci. 127, 3217– 3225 (2014).

59. Vuoristo, S. et al. Laminin isoforms in human embryonic stem cells: synthesis, receptor usage and growth support. J. Cell. Mol. Med. 13, 2622–2633 (2009).

60. Sauvanet, C., Wayt, J., Pelaseyed, T. & Bretscher, A. Structure, Regulation, and Functional Diversity of Microvilli on the Apical Domain of Epithelial Cells. Annu. Rev. Cell Dev. Biol. 31, 593–621 (2015).

61. Niazi, A. et al. Microvilli control the morphogenesis of the tectorial membrane extracellular matrix. Dev. Cell 60, 679–695.e8 (2025).

62. Fattah, A. R. A. et al. Extracellular matrix flow guides in-vitro epithelial morphogenesis. 2022.10.21.513217 Preprint at 10.1101/2022.10.21.513217 (2022).

63. Bhave, G., Colon, S. & Ferrell, N. The sulfilimine cross-link of collagen IV contributes to kidney tubular basement membrane stiffness. Am. J. Physiol. Renal Physiol. 313, F596–F602 (2017).

64. Khalilgharibi, N. & Mao, Y. To form and function: on the role of basement membrane mechanics in tissue development, homeostasis and disease. Open Biol. 11, 200360 (2021).

65. Bonnans, C., Chou, J. & Werb, Z. Remodelling the extracellular matrix in development and disease. Nat. Rev. Mol. Cell Biol. 15, 786–801 (2014).

66. Lampi, M. C. & Reinhart-King, C. A. Targeting extracellular matrix stiffness to attenuate disease: From molecular mechanisms to clinical trials. Sci. Transl. Med. 10, eaao0475 (2018).

67. Llewellyn, J., Charrier, A., Cuciniello, R., Helfer, E. & Dono, R. Substrate stiffness alters layer architecture and biophysics of human induced pluripotent stem cells to modulate their differentiation potential. iScience 27, 110557 (2024).

68. Vasic, I. et al. Loss of TJP1 disrupts gastrulation patterning and increases differentiation toward the germ cell lineage in human pluripotent stem cells. Dev. Cell 58, 1477–1488.e5 (2023).

69. Legier, T. et al. Epithelial disruption drives mesendoderm differentiation in human pluripotent stem cells by enabling TGF-β protein sensing. Nat. Commun. 14, 349 (2023).

70. Fort, L., Vamadevan, V., Wang, W. & Macara, I. G. Actomyosin Contractility is a Potent Suppressor of Mesoderm Induction by Human Pluripotent Stem Cells. 2024.09.30.615859 Preprint at 10.1101/2024.09.30.615859 (2025).

71. Levental, K. R. et al. Matrix crosslinking forces tumor progression by enhancing integrin signaling. Cell **139**, 891–906 (2009).

72. Fiore, V. F. et al. Mechanics of a multilayer epithelium instruct tumour architecture and function. Nature 585, 433–439 (2020).

73. Stringer, C., Wang, T., Michaelos, M. & Pachitariu, M. Cellpose: a generalist algorithm for cellular segmentation. Nat. Methods 18, 100–106 (2021).

